# Direct genome sequencing of *Leishmania tropica* in tissues of Moroccan patients with cutaneous leishmaniasis reveals micro-focal transmission underlain by clonal and sexual reproduction modes

**DOI:** 10.1101/2025.04.17.649263

**Authors:** Othmane Daoui, Pieter Monsieurs, Hasnaa Talimi, Gerald F. Späth, Jean-Claude Dujardin, Senne Heeren, Meryem Lemrani, Malgorzata Anna Domagalska

## Abstract

*Leishmania tropica* is causing cutaneous leishmaniasis (CL) from North Africa to India and in Ethiopia and is reported to be transmitted from humans to humans through sand fly bites. While this species is characterized by a high genetic diversity all over the area of endemicity, there is very little information on diversity at micro-epidemiological scale. Here, we zoomed on an epidemic Moroccan focus of CL - restricted in space and time - and studied transmission patterns by comparative genomics of parasites in human patients. We used a culture-independent method of genome sequencing based on *Leishmania* genome capture (SureSelect-sequencing, SuSL-seq), applied directly on dermal scrapings. We first compared the genome of paired samples: i.e. parasites in host tissues analyzed by SuSL-seq and derived isolates shortly maintained in culture and analyzed by whole genome sequencing (WGS). Despite the low passage number of isolates, significant differences were observed between karyotypes of 4/7 paired samples, highlighting the clinical and epidemiological relevance of direct genome sequencing. Secondly, we identified 7 groups of nearly identical genotypes, characteristic of clonal propagation as well as parasites with mixed ancestry, a signature of genetic exchange. Our results reveal a micro-focal transmission among humans, underlain by clonal and sexual reproductive modes. This study demonstrates the power of direct genome sequencing for evolutionary genetics at a micro-epidemiological scale.

## Introduction

Leishmaniasis is a complex of vector borne parasitic diseases of major public health importance: visceral leishmaniasis (VL: 13,814 new cases in 2019 and 510,000 DALYs [Disability-adjusted life years] in 2017) is lethal in the absence of treatment and cutaneous leishmaniasis (CL: 277,224 new cases in 2019 and 260,000 DALYs in 2017) causes serious disability, stigmatization and mental health problems [1]. The disease is (re-)emerging worldwide owing in part to anthropogenic and environmental changes, global mobility, drug resistance and immuno-suppression [2]. A few control programs have been launched, like the (so far) successful VL elimination program in the Indian sub-continent (ISC) [3], but their long-term efficacy is not yet guaranteed: (i) in a post-elimination phase, where political support and means may decrease, (ii) history revealed the cyclical nature of epidemics with long inter-epidemic periods when the parasites are hardly detected and (iii) intervention tools like drugs or diagnostics may lose efficacy. In contexts like these, continuous surveillance is essential, but it is striking how genomic surveillance remains neglected for parasites like *Leishmania*, despite demonstrating its value for monitoring the evolution and spreading of other pathogens, like viruses during the COVID-19 crisis [4]. Yet, genomic surveillance of *Leishmania* theoretically allows answering several questions of major public health importance [5]. Illustrating this broad portfolio with specific and practical studies should contribute to counter this neglect and justify the implementation of genomic surveillance.

Thanks to its highest discriminatory power, whole genome sequencing (WGS) might become a gold standard for *Leishmania* genotyping. The power to answer clinical and epidemiological questions with WGS was already demonstrated for some of them, (i) we traced the evolution of *L. donovani* and VL epidemics in the Indian sub-continent (ISC) over the last 150 years [6]; (ii) we identified the source of a recent VL outbreak in Nepal [7]; (iii) others found novel and less virulent *L donovani* strains in Sri Lanka, arising through the introgression of *L. major* or *L. tropica* genomes in *L. donovani* [8]; (iv) we detected an unprecedented level of sexual recombination in *L. braziliensis* from Southern Peru and established the probable role of hybridization in the spreading of its endogenous double stranded RNA virus (*Leishmaniavirus*; LRV1) and treatment failure [9]. However, to our knowledge, no genomic study was so far dedicated to the characterization of *Leishmania* transmission at a micro-epidemiological scale.

*Leishmania tropica* is a species causing CL and it is endemic essentially in North Africa, Middle East and India/Pakistan, albeit an outbreak was recently reported in Ethiopia [10,11]. The species is reported to be transmitted by sand flies (Phlebotominae) and mostly from humans to humans, but some studies mention a possible contribution of hyraxes, bats and wild rodents in transmission [12,13]. *Leishmania tropica* was reported to be the most polymorphic *Leishmania* species complex in the Old World [14]. Recently, the genomic diversity of *L. tropica* in whole Morocco was analyzed and compared to other endemic countries [15]. This revealed genetic heterogeneity within Morocco and allowed to differentiate Moroccan *L. tropica* strains from those circulating in the Middle East. Here, we zoomed in an epidemic focus - restricted in space and time- and looked for micro-focal transmission patterns by comparative genomics of the parasites in CL patients. A major conceptual difference with previous studies - all directed to cultivated isolates-is that we applied here a method of direct genome sequencing in human host tissues, based on *Leishmania* genome capture with a set of 194,462 parasite-specific RNA probes (SureSelect-sequencing, SuSL-seq). The method was developed and validated in *L. donovani* [7,16] and provides two major advantages in comparison to approaches targeting cultivated isolates [5]. On one hand, it allows a broader representation of the parasite’s genomic diversity, given its independence from isolation success rate which varies from one laboratory to the other and is far from perfect. On the other hand, it is not biased by the isolation bottleneck and genomic adaptation occurring during in vitro cultivation [17], hence it is *a priori* more adequate to understand the biology and the epidemiology of the parasites in their natural environment, i.e. macrophages in the vertebrate hosts, than in cultivated life stages.

In this study, we first compared the genome of paired samples (parasites in biopsies and derived isolates analyzed by SuSL-seq and WGS respectively) available from some of the patients of Foum Jemaa, in order to assess the patterns of in vitro genomic adaptation in *L. tropica*. Secondly, we applied a phylogenomic approach to study the recent evolution of *L. tropica* population in the focus and transmission patterns in a spatial context.

## Methods

### Ethics, patients and samples

Clinical samples were collected from CL patients recruited in the city of Foum Jemaa. Study was approved by the Ethical Review Committee for Biomedical Research of the Faculty of Medicine and Pharmacy, Rabat, Morocco (IORG 0006594 FWA00024287) and the Institutional Review Board of the Institute of Tropical Medicine of Antwerp (ITMA), Belgium (file 1561/22). All adult participants provided informed consent, including for secondary use of data and samples. For the inclusion of young children, consent was obtained from the respective parents of legal guardians. From each patient, a dermal scraping sample was collected and DNA extracted for molecular analyses, with the Qiagen kit. DNA from 17 qPCR-positive samples for *Leishmania* was studied here.

Details on the samples here studied are indicated in Table S1, Fig.S1 and Fig.1. Seventeen clinical samples (prefix SuSL) originated from Foum Jemaa (N=14) and Imintanout (N=3) were sequenced in the present study. For 7 of the clinical samples, paired cultivated isolates were available and these have been sequenced elsewhere [15]; in addition, the previously sequenced [15] genomes of 7 Moroccan isolates for which there was no paired clinical samples were added as reference/outgroups.

**Fig. 1.**
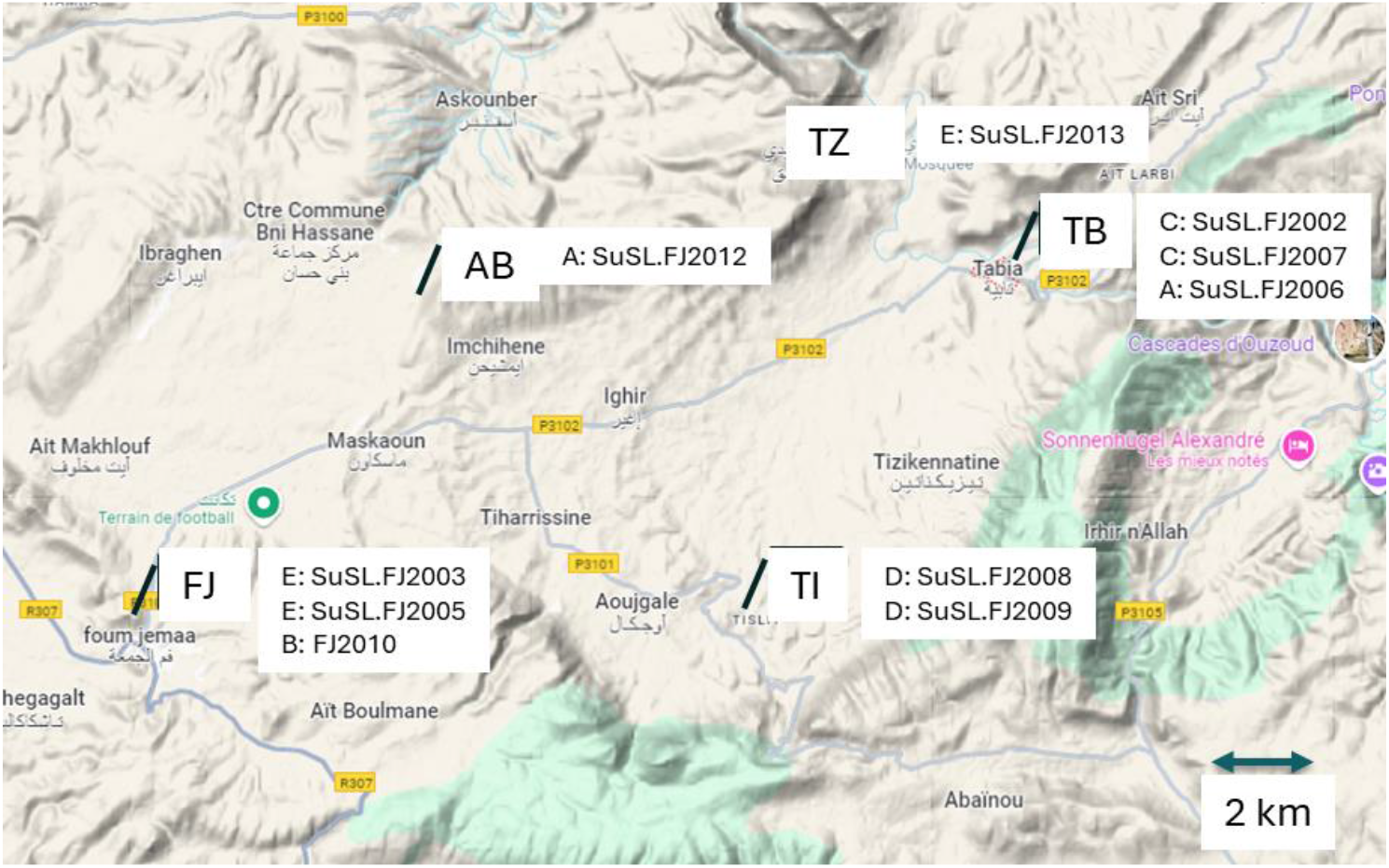
Map of Foum Jemaa focus, with geo-localization of the origin of 10 clinical samples for which coordinates of the locality of origin were available (Foum Jemaa itself, FJ; Tabia, TB; Tislit, TI; Ait Bouchao, AB and Tiazite, TZ,); if not, samples were allocated to ‘Foum Jemaa focus’, but not indicated on the map. The codes of the 7 groups of nearly-identical genotypes (A-G, see Fig.3) are indicated; no code means that the sample branched outside the groups. See also Fig.3. Map: © 2024 Google.

### Processing of samples for sequencing

For each sample, DNA was extracted using the QIAamp DNA Blood Mini Kit (Qiagen, Venlo, the Netherlands) following the manufacturer’s instructions. *Leishmania* species identification was performed by ITS1-PCR followed by enzymatic restriction by HaeIII [18]. For the current study, we analyzed 17 samples from patients infected with *L. tropica*. The use of SureSelect technology requires (i) a minimum DNA input at least 10 ng, which can be difficult to reach when working with biological samples and (ii) a % Leish DNA above 0.006% [16]. The two parameters were assessed here. On one hand, DNA was quantified with the Qubit fluorometric assay and 11 clinical samples underwent DNA concentration using AMPure beads as done elsewhere [11]. On the other hand, the percentage of *Leishmania* (vs human) DNA was calculated by qPCR as reported elsewhere [16]. The 17 samples showed a % Leish DNA above 0.047 % (median, 2.31 %, Table S1). The 17 samples (11 concentrated and 6 not concentrated) were then submitted to target enrichment. SureSelect (Agilent Technologies) was used to capture *Leishmania* genomic DNA following standard SureSelect XTHS Target Enrichment system protocol for Illumina Multiplexed Sequencing platforms using either the *L. donovani / L. infantum* design (Agilent SureSelect design ID S3377046) or the *L. aethiopica* design (design ID S3373475).

### Bio-informatic analyses

In addition to the sequencing data produced within this work, we also used sequencing data of *L. tropica* strains obtained within Morocco ([15], BioProject accession number PRJNA978932 and PRJNA422027). All sequencing data were processed as described elsewhere [7]. In summary, reads were aligned to the *L. tropica* L590 reference genome (TriTrypDB release 64) using BWA (v0.7.17), with a seed length of 50. Only properly paired reads with a mapping quality >30 were retained using SAMtools. Duplicate reads were removed with Picard (v2.22.4) using the RemoveDuplicates function. SNP calling followed the GATK (v4.1.4.1) best practices workflow: (1) HaplotypeCaller was used to generate GVCF files for each sample; (2) individual GVCFs were merged with CombineGVCFs; (3) genotyping was performed with GenotypeGVCFs; and (4) SNP and INDEL filtering was applied using SelectVariants and VariantFiltration. Variant regions corresponding to known drug resistance markers were extracted using BCFtools and visualized with R’s pheatmap package. Overlap of SNPs between two strains was calculated using the vcfR (version 1.13.0) and adegenet (version 2.1.8) package in R, and visualization was done using the pheatmap (version 1.0.12) package.

Phylogenetic analysis was conducted with RAxML. Bi-allelic SNPs were extracted from VCF files using BCFtools and converted to Phylip format with vcf2phylip.py Python script https://github.com/edgardomortiz/vcf2phylip. Phylogenetic trees were constructed under the GTR+G model with 1000 bootstrap replicates, using *L. aethiopica* L147 as an outgroup. Tree visualization was performed with ggtree. For the unrooted phylogenetic networks, biallelic SNPs were converted to a fasta file using the vcf2fasta.py Python script (https://github.com/FreBio/mytools/tree/master) and visualized using SplitsTree4 with the NeighborNet algorithm [19].

For downstream population genomic analysis, bed files were generated using PLINK 2.0 after pruning SNPs for sites with high linkage dequilibrium (--indep-pairwise 50 10 0.5). ADMIXTURE was employed to assess the ancestry of both WGS and SureSelect samples, excluding clonal individuals within each phylogenetic group. The number of populations (K) was tested from 2 to 4, determined through a five-fold cross-validation procedure. Population structure inferred by ADMIXTURE was visualized using ggplot2 in R.

Two reference genomes of *L. tropica* are available: the TriTrypDB (release 64) *L. tropica* L590 and the NCBI (strain CDC216-162) reference. The latter one appeared not to be a bona fide *L. tropica*. Accordingly, for the analysis of genome nucleotide sequences, we mapped reads of *L. tropica* samples to the TriTrypDB reference. For the analysis of aneuploidy, there was a problem with somy of chr31 of the TriTryp reference and we used the reference genome of *L. aethiopica* L147 as available on TriTrypDB (release 64), the species phylogenetically closest to *L. tropica* [20]. This is explained in detail in the Supplementary results (Figs S2-S5).

## Results

### 1. Comparison of different SuSL-seq designs for capture and sequencing of *L. tropica* genome

In order to optimize cost-efficacy of direct sequencing and avoid the need to develop specific SuSL designs for any new species to be studied, we tested on *L. tropica* experimental samples two SuSL-designs that were available in the laboratory, respectively targeting *L. donovani* and *L. aethiopica*. The rationale was that SuSL probes should hybridize with regions of higher similarity, even better in phylogenetically related species: we therefore expected here that *L. aethiopica* probes would provide better results on *L. tropica* than *L. donovani* [20], which was verified (see supplementary results, Fig.S6a). Consequently, *L. aethiopica* array was applied to the 17 clinical samples: in 15 of them the % of reads mapping to the TriTrypDB *L. tropica* reference genome was above 75%, 20-40 % for the 2 remaining ones (Fig.S6b). We pursued our analyses with the 17 samples.

### 2. Comparison between direct sequencing of clinical samples with SuSL-seq and sequencing of cultivated isolates with WGS

In the phylogenomic analyses described here below, two types of samples were combined, i.e. cultivated isolates and clinical samples. In previous work on *L. donovani*, we reported genomic plasticity between the respective parasites. Here, we verified and assessed the extent of the phenomenon, using paired samples (dermal scrapings and derived isolates) from 7 CL patients. Corresponding genome sequences (SuSL-seq for dermal scrapings and WGS for isolates) were compared for 4 elements of genomic variation: SNPs, INDELs, local copy number variations (CNVs) and chromosome copy number variation. For local CNVs, there appeared to occur a systematic technical bias between copy number in the host and derived isolate, hence CNVs were excluded from further analyses (see Supplementary results: Fig.S7 and S8). We thus further focused on the 3 remaining diversity elements (see summary in Table 1): SNPs, INDELS and chromosome copy number variation.

**Table 1.**
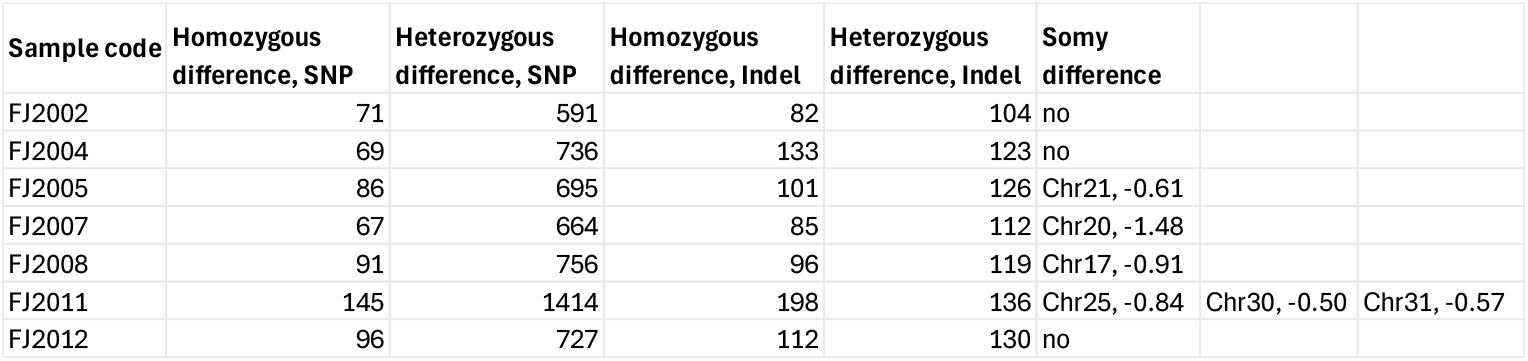
Genomic differences observed between paired samples (dermal scrapings from 7 patients and respective 7 derived isolates. Karyotype: S is the S-value (average somy for a given chromosome); we mention only if a given chromosome (chr) shows a significant difference in somy (i) S value of SuSL-seq sample – S value of WGS sample being ≥ 0.5 and (ii) p value < 0.05.

| Sample code | Homozygous difference, SNP | Heterozygous difference, SNP | Homozygous difference, Indel | Heterozygous difference, Indel | Somy difference |  |  |
| --- | --- | --- | --- | --- | --- | --- | --- |
| FJ2002 | 71 | 591 | 82 | 104 | no |  |  |
| FJ2004 | 69 | 736 | 133 | 123 | no |  |  |
| FJ2005 | 86 | 695 | 101 | 126 | Chr21, -0.61 |  |  |
| FJ2007 | 67 | 664 | 85 | 112 | Chr20, -1.48 |  |  |
| FJ2008 | 91 | 756 | 96 | 119 | Chr17, -0.91 |  |  |
| FJ2011 | 145 | 1414 | 198 | 136 | Chr25, -0.84 | Chr30, -0.50 | Chr31, -0.57 |
| FJ2012 | 96 | 727 | 112 | 130 | no |  |  |

Overall, there was a relatively small number of SNP differences between clinical samples and derived isolates, ranging from 67 to 145 homozygous differences and from 591 to 1414 heterozygous differences. For INDELs, the number of homozygous differences ranged from 82 to 198 and the number of heterozygous differences was in the same range (from 104 to 136). Comparison of karyotypes using the Student’s t-test revealed a significant difference in 4/7 patients and concerned a total of 6 chromosomes. In all cases, the somy value increased from SuSL-seq to WGS sample (Table 1, Table S2 and Fig. S9). In all SuSL-seq samples, chromosomes were disomic with the exception of Chr31 which is always reported to be tetrasomic. Noteworthy, in one patient (FJ20_08) Chr17 was monosomic, in the SuSL-seq samples, while it was disomic in the derived isolate (Fig.2).

**Fig. 2.**
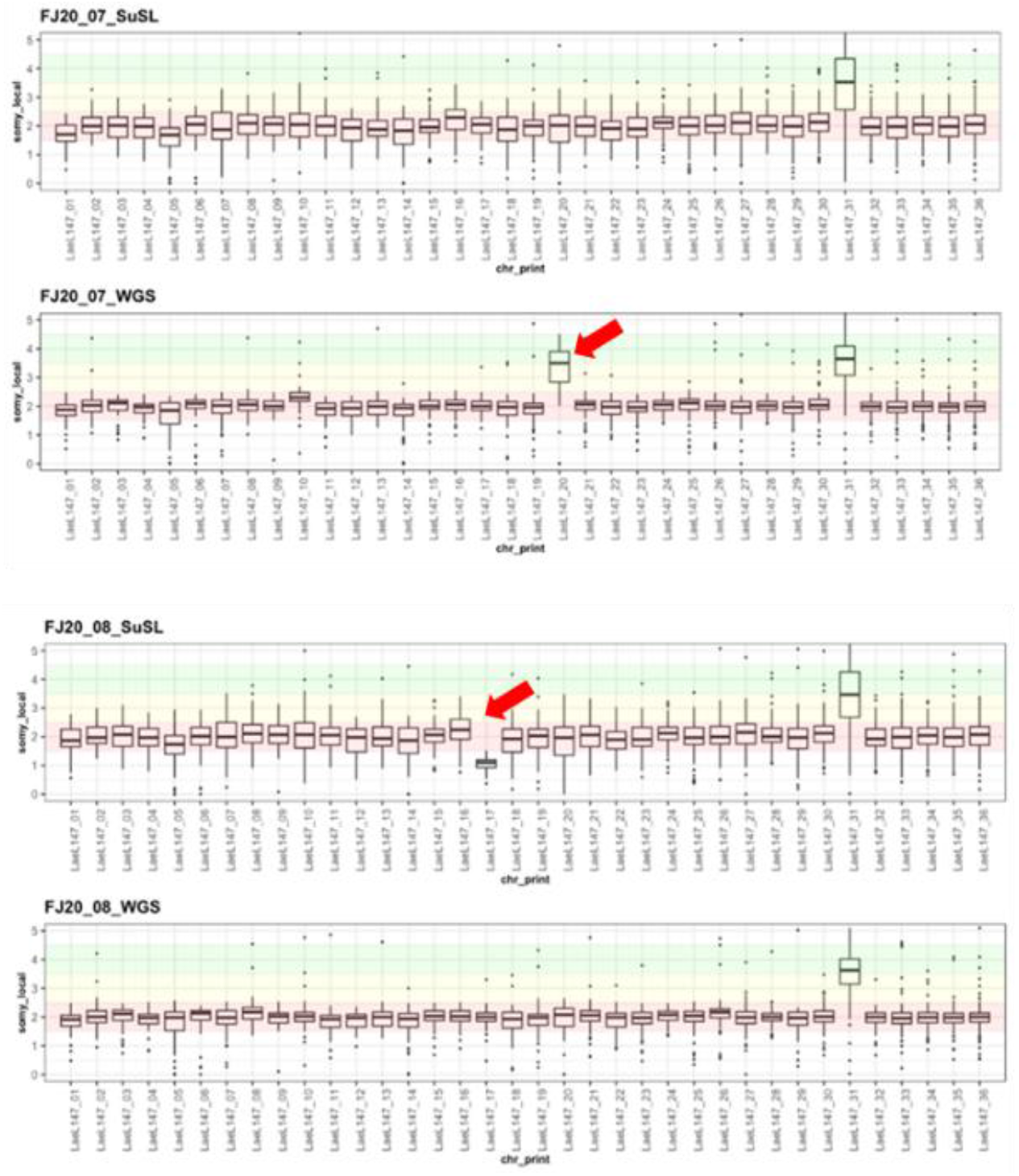
Karyotypes of paired samples (for each patient, SuSL-seq, clinical sample and WGS, derived isolate); the arrow marks the chromosomes showing a significant change in somy between paired samples (see criteria in the legend of Table 1; 2 representative cases are shown here, see Fig.S9 for complete analysis of 7 paired samples.

### 3. Phylogenomic analyses

We first did an analysis of all samples and isolates included in this study, in order to situate the parasite population of the Foum Jemaa focus in a broader Moroccan context (Fig.S1). Inspection of Fig.S10 shows that the phylogenetically most remote parasites (M3015 and SuSL.IT02, −04 & −08) were also geographically the most distant (respectively Ouarzazate - at the other side of the Atlas Mountains- and Imintanout at more than 200 km). Within the triangle constituted by Foum Jemaa focus, Tannant and Azilal (side, max 50 km), phylogenetic distances were relatively much smaller. Two clinical samples from Imintanout (IT02 and IT04) clustered closely together, constituting a group of nearly identical genomes (254 homozygous SNPs, see below)

In a next step, we removed the phylogenetically and geographically distant parasites (OU and IM) and zoomed on the samples from the triangle Foum Jemaa-Tannant-Azilal, with an emphasis on Foum Jemaa focus. We first applied hierarchical clustering (Fig.S11) and identified 7 groups of nearly identical genomes (here defined as genomes showing less than 500 homozygous SNPs differences with other genomes of the same group; blue boxes in Fig.S11), named A to G. Secondly, we performed a phylogenetic analysis, identified the 7 groups and geo-localized samples, when possible (Fig.3). This revealed 3 findings:

**Fig. 3.**
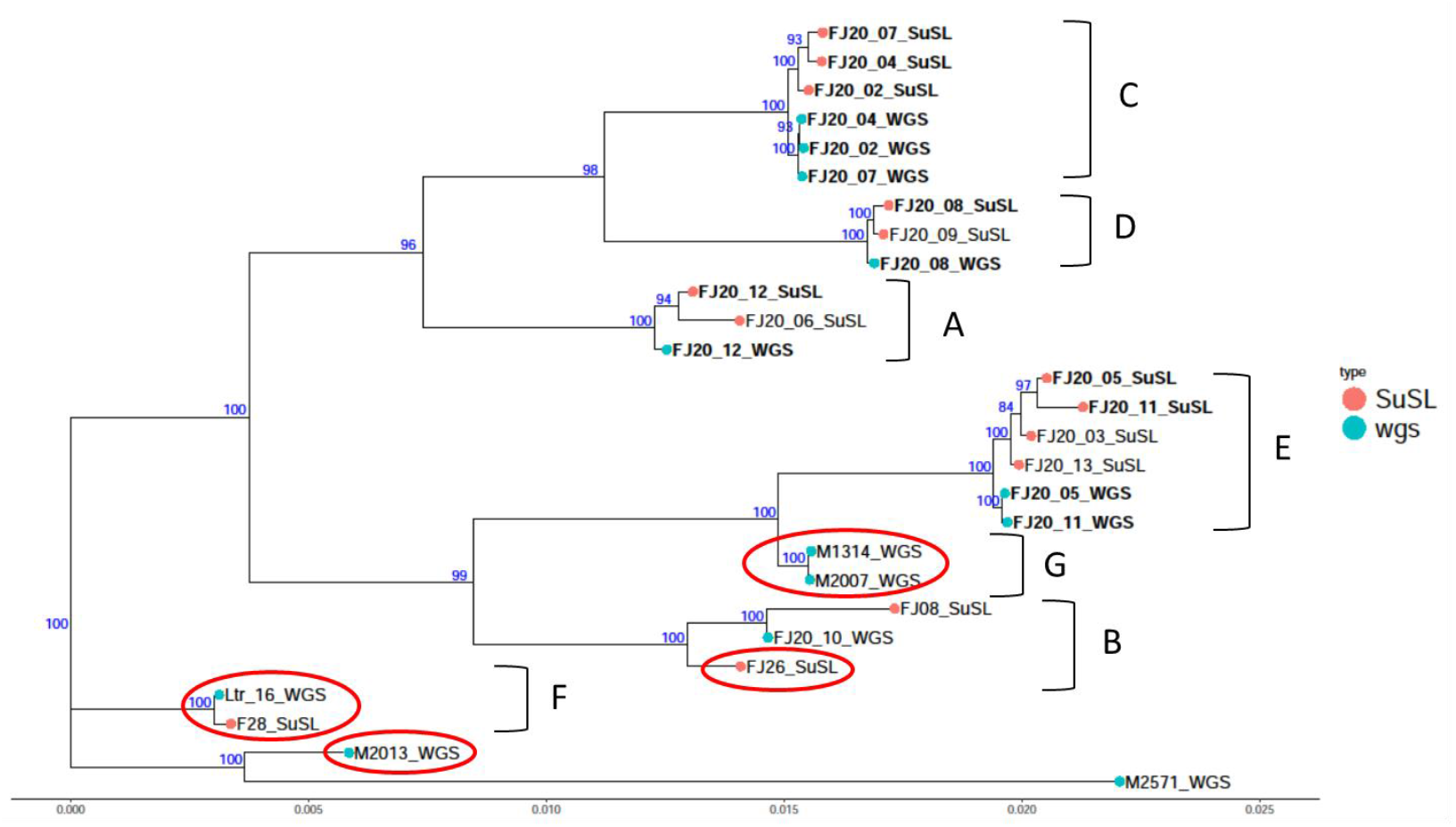
Rooted phylogenetic tree based on genome-wide bi-allelic SNPs using RAxML, with an indication of the bootstrap values in blue; A-G are the 7 clusters of nearly-identical genomes observed in Supp Fig.5; All samples with FJ prefix come from the ‘Foum Jamaa focus’ (see M&M). The six samples highlighted in red correspond to samples with mixed ancestry, as revealed by admixture analysis (see Fig. 4); the seventh admixed sample (M3015) was an outlier in phylogenetic analysis (Fig.S10) and is thus not represented here.

1. for each of the 7 paired samples, genomes of clinical samples and derived isolates clustered together in the same group (SuSL/FJ2002, FJ2004 and FJ2007 in group C; SuSL/FJ2008 in group D; SuSL/FJ2012 in group A; SuSL/FJ2005 and FJ2011 in group E).
2. in 3 localities, 2 clinical samples of the group of nearly identical genotypes were observed in different patients: FJ20_03 and FJ20_05 (E) in Foum Jemaa, FJ20_08 and FJ20_09 (D) in Tislit and FJ20_02 and FJ20_07 (C) in Tabia. This suggested local transmission in each of the 3 localities but did not exclude the presence of additional and different genotypes in the localities: e.g. FJ20_10 together with 2 samples from group E in Foum Jemaa or FJ20_06 together with 2 samples from group C in Tabia.
3. in addition, two genetic groups were present in two distinct localities: group E (in Foum Jemaa and Tiazite; ca. 20km apart) and group A (in Ait Bouchao and Tabia; ca. 19 km apart).

### 4. Tracing signatures of recombination

The sympatric sample analyzed in the present study together with the high discriminatory power of whole genome sequencing also provided unique conditions to search for possible signatures of recombination. First, a phylogenetic network was made with the NeighborNet algorithm. This revealed a reticulate network, which may suggest the occurrence of recombination (Fig. S12). In a next step, analysis of population structure was done with ADMIXTURE based on 8474 genome-wide bi-allelic SNPs after Linkage Disequilibrium pruning. Line plot shows the cross-validation error (CV) for K = 3 to K = 10 (lowest CV-error: K = 4; Fig. S13). Most samples showed a single ancestral origin (Fig. 4). In contrast, for K=4 and K=5, seven (5 + 2 nearly identical genotypes) samples showed a clear mixed ancestry (Fig. 4).

**Fig. 4.**
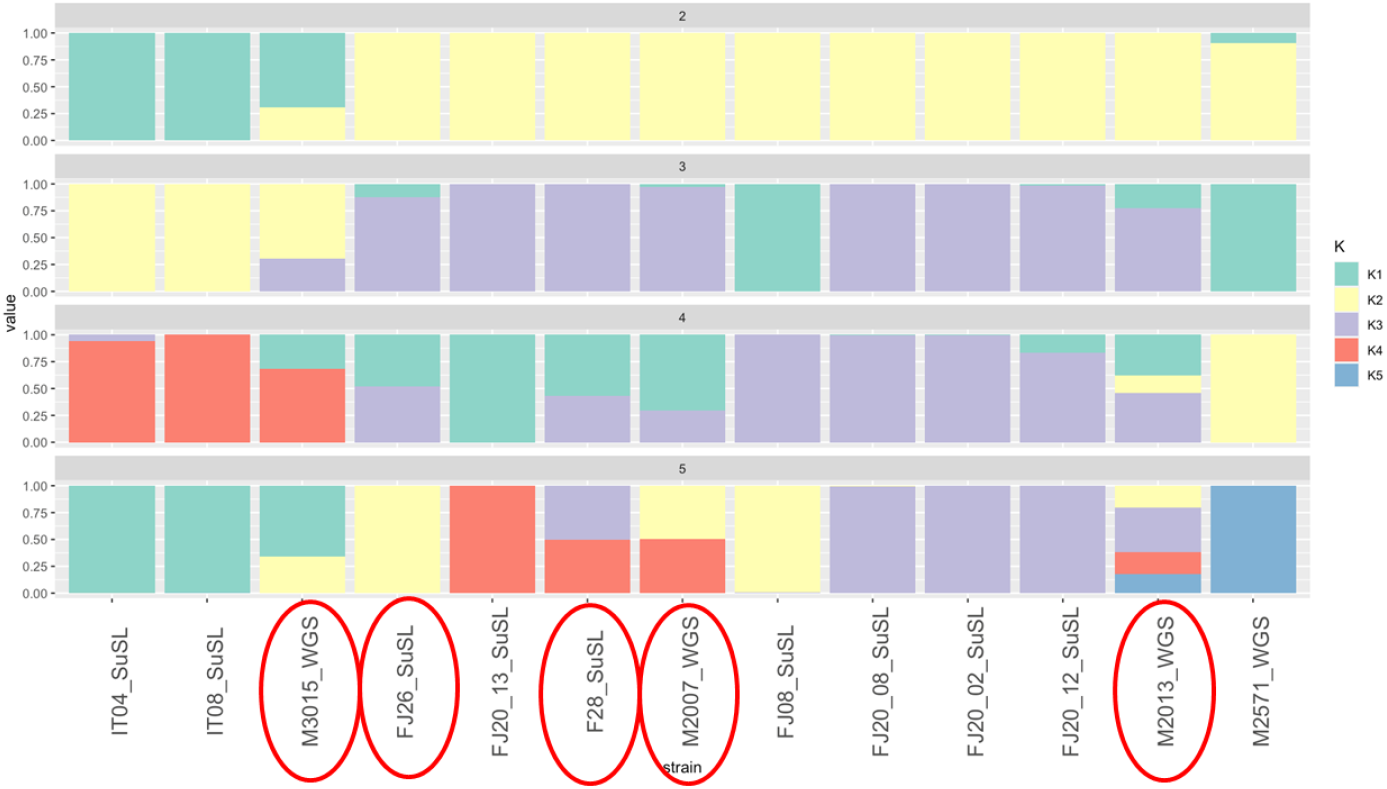
ADMIXTURE analysis showing ancestry proportions of *L. tropica* samples across population clusters (K = 1 to K = 5). Each bar represents a sample, with colors corresponding to ancestry components. Samples with mixed ancestry are highlighted in red ovals. Nearly identical genotypes, such as Ltr16 (F28) and M1314 (M2007), were excluded but are counted as admixed, bringing the total number of admixed samples to seven.

Presence of mixed ancestry signatures in a sample could be reflecting events of genetic exchange as well as polyclonality (mixture of different genotypes). To distinguish the two hypotheses, we plotted the alternative allele frequencies along the 36 chromosomes of admixed samples. In case of polyclonal mixture, alternative allele frequency should be constant all along the chromosomes and it should deviate from 50/50. While in case of genetic exchange, stretches of heterozygosity should be alternating with stretches of homozygosity (intra-chromosomal recombination) and in heterozygous stretches, the frequency should always be 50/50 [21]. The latter was observed for most chromosomes of admixed samples (see example for chromosome 27, Fig.5). Interestingly, samples with single ancestral origin also showed alternating stretches of homozygosity and heterozygosity, suggesting more intra-chromosomal recombination than originally suggested by admixture analysis.

**Fig. 5.**
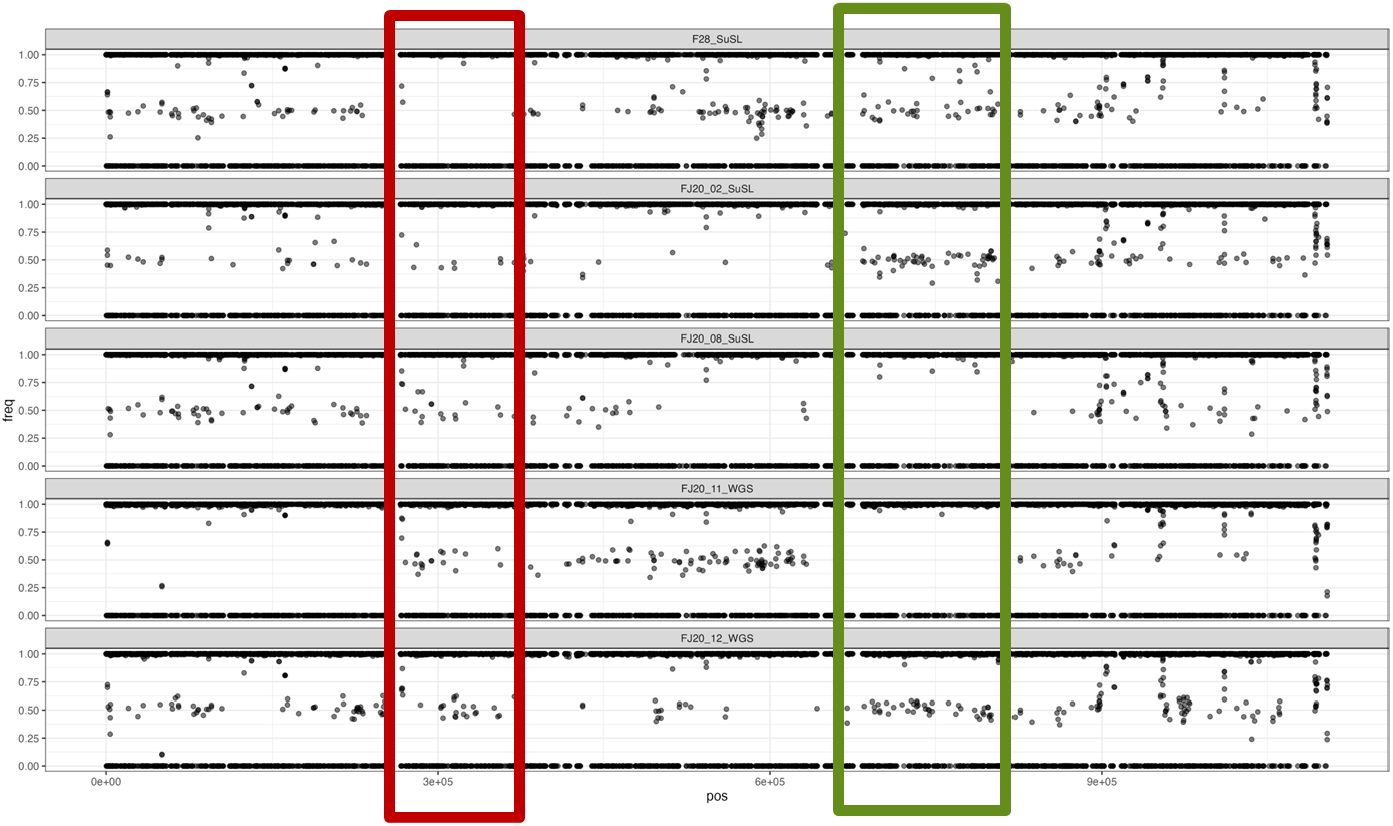
Allele frequency plot for chromosome 27 for five different samples. The X-axis represents the position of the SNP in the chromosome, and the Y-axis represents the allele frequency of the SNP in the corresponding strain. The red box highlights a homozygous region in F28_SuSL, while the green box highlights a homozygous region in strains FJ20_08_SuSL and FJ20_11_WGS.

## Discussion

In North Africa, CL is highly prevalent in Morocco, Algeria, Tunisia, and Libya. It is due to 3 species, two strictly dermotropic i.e. causing CL (*L. tropica* and *L. major*) and one causing both CL and VL (*L. infantum*). The eco-epidemiology of the three species is reported to be different: on one hand, *L. major* and *L. infantum* are both zoonotic, with wild rodents and canids as respective reservoir; on the other hand, *L. tropica* is considered to be transmitted in an anthroponotic way [22], even if a zoonotic spillover cannot be excluded [23]. In Europe, CL due to *L. tropica* is an imported pathology, encountered essentially among migrants (80%) [24]. In 2010, the risk of introducing exotic *Leishmania* species in Europe was considered to be low, because of the absence of proven vectors and/or reservoir hosts [25]. However, more recently the danger of local transmission of *L. tropica* was evidenced in Greece [26] and in Spain [27]. Molecular surveillance is clearly needed, to understand the transmission and circulation of the parasite at global and local levels and this for better patient management and control of the disease.

Early studies of the molecular epidemiology of leishmaniasis in Morocco were done with targeted methods such as multi-locus enzyme electrophoresis (MLEE [14]) or Multilocus sequence analysis (MLST [28]). MLEE allowed to describe a large enzymatic diversity in 1988-90 in the Azilal province (Foum Jemaa, Tanant, here studied too), with the observation of 7 zymodemes (groups of parasites characterized by a same MLEE profile). This diversity was interpreted to be a consequence of the ‘old’ age of the focus (allowing the colonization by different genetic groups of parasites) as well as a state of hypo-endemic equilibrium at the time of sampling (while a single zymodeme would be the signature of hyper-endemicity or outbreak) [14]. A similar level of diversity was shown by a MLST study conducted in 2018: the sequence of 5 parasite’s genes in 13 human samples from Foum Jemaa revealed 8 haplotypes [28]. Even if these two studies and the present one were done on samples from the same region, it is difficult to compare them, as sampling was done at different periods with molecular methods showing major differences in discriminatory power. However, the overall genetic structure here provided by SuSL-Seq and WGS fits with that observed with MLEE and MLST, with 7 genomic groups each differing by 1154 SNPs, while the average nucleotide dissimilarity within each group was lower than 500. In comparison, in a recent study on *L. tropica* outbreak in Ethiopia, we found only nearly identical genotypes, with an average of 122 nucleotide differences between them [11]. Thus, following the classification by Pratlong et al. [14], the picture revealed by WGS would suggest a combination between hypo-endemic equilibrium (between 7 genetic groups) and micro-epidemic expansion within each of these groups.

Two *Leishmania* reproductive modes underly this complex epidemiological pattern. On one hand, micro-epidemic expansion fits with the clonal propagation here revealed by (i) the nearly identical genotypes each observed within 3 localities and (ii) the pure ancestry observed for most samples in admixture analysis. On the other hand, the same admixture analysis showed a mixed ancestry in 5 samples; this, together with the topology of the phylogenetic network and the long branch separating these 5 samples from the other ones might constitute a possible signature of genetic exchange in 5 samples; this would have occurred at least three times: first, generating F28-SuSL and Ltr16_WGS, secondly, generating M1314_WGS and M2007_WGS and thirdly, generating FJ26_SuSL. Previous studies have revealed naturally occurring sexual recombination involving *L. tropica* and other species, like *L. aethiopica* [29] or *L. donovani* [8], but also occurring at intra-species level [30]

The high discriminatory power and the capacity to be used directly on host’s tissues, with high genome coverage (more than 70% in most samples here studied) are among the greatest advantages of of SuSL-seq. This power was used in a previous study for source tracing of *L. donovani* in emerging foci of VL in Western Nepal [7] and for describing a new emerging focus of CL in Ethiopia [11]. In the present study, the precise geo-localization of a sub-set of samples provided information about the micro-focal patterns of transmission in the region. This showed that groups A, B and C were encountered in Foum Jemaa, Tabia and Tislit respectively, each time with 2 independent samples, suggesting local transmission. Furthermore, one sample of group A was geo-localized in Tiazite (20 km from FJ). It is impossible at this stage to know if the CL patient from Tiazite got infected during a stay in Foum Jemaa or if group A parasites spread to Tiazite and were also transmitted in that locality. This clearly indicates that human mobility plays a major role in the dynamics of transmission. Further studies should require a larger sample, include insect vector and re-visit the possibility of animal reservoir, including synanthropic animals.

In a previous report, we analyzed paired bone marrow samples of VL patients and derived *L. donovani* isolates and compared the genome of the respective parasites (applying SuSL-seq for amastigotes in bone marrow and WGS for isolated promastigotes) and found significant differences, especially at karyotype level [16]. In amastigotes, the majority of chromosomes showed to be disomic and when they were put in culture, the parasites developed rapidly aneuploidy, likely of adaptive significance [17,31]. A similar observation was made here with *L. tropica* paired samples. Even if an attempt was made to limit aneuploidy changes in cultivated promastigotes by keeping low (≤ 5) the number of passages since isolation, some chromosomes already showed some changes in somy in isolates and the monosomy of chr17 in amastigotes of SuSL-FJ2008 disappeared in derived isolate, where the chromosome was disomic. The functional significance of this monosomy is unknown: so far, we observed this phenomenon essentially in clinical samples.

The present study demonstrates the power of SuSL-seq to study *Leishmania* transmission patterns and evolutionary genetics at a micro-epidemiological scale, and this without the need for parasite isolation and cultivation. Our method offers the sensitivity and discriminatory power to genotype the parasites in sand flies and animal reservoir (if any) and (re-)visit the circulation of the parasite throughout the whole life cycle. This can be highly relevant for studies of outbreaks, identification of risk factors, detection of genomic signatures of drug resistance, surveillance and monitoring of control interventions. From a genetic point of view, our approach also offers the resolution needed to detect sexual reproduction at an intra-species level, between closely related parasites. Further work would be needed to re-visit the balance between clonal and sexual reproduction modes in real-life and understand its clinical and epidemiological impact.

## Notes

## Data availability

All sequencing data are available at http://www.ncbi.nlm.nih.gov/bioproject/1246653 (PRJNA1246653). All other data are available in the main text or supplementary material.

## Acknowledgments

This study was supported by the EU-funded project LeiSHield-MATI RISE ‘A multi-disciplinary international effort to identify clinical, molecular and social factors impacting cutaneous leishmaniasis’ (Grant Agreement No. 778298). SH was supported by the Research Foundation Flanders (grants G092921N); MAD acknowledges support from Flemish Ministry of Science and Innovation.

## AI

The authors did not use artificial intelligence to write this manuscript

