## supplementary material for "Direct genome sequencing of *Leishmania tropica* in tissues of Moroccan patients with cutaneous leishmaniasis reveals micro-focal transmission underlain by clonal and sexual reproduction modes"

[1]Laboratory of Parasitology and Vector-Borne-Diseases, Institut Pasteur du Maroc, Casablanca, Morocco; [2]Molecular Parasitology Unit, Institute of Tropical Medicine, Antwerp, Belgium; [3]Experimental Parasitology Unit, Institute of Tropical Medicine, Antwerp, Belgium; [4]Systems and Data Engineering Team, National School of Applied Sciences, University Abdelmalek Essaadi, Tangier, Morocco ; [5]Unité de Parasitologie Moléculaire et Signalisation, Institut Pasteur, Université Paris Cité, INSERM 1201, , Paris, France. [6]Department of Microbiology, Immunology and Transplantation, Rega Institute for Medical Research, Katholieke Universiteit Leuven, Leuven, Belgium.

### Equal contribution

* Co-senior and corresponding authors

Meryem Lemrani, 1 Place Louis Pasteur, Casablanca 20360, Morocco

Malgorzata Anna Domagalska, 155 Nationalestraat, Antwerpen, 2000, Belgium

#### 1. Samples

**Table S1**. List of samples: for cultivated isolates, WHO code is used and their genome sequence was reported elsewhere [1]. Clinical sample codes were pseudonymized, and only the clinicians were able to link the patient names to the study codes. The Foum Jemaa focus corresponds to 5 entities (Foum Jemaa itself, FJ and neighborhoods or surrounding localities: Tabia, TB; Tislit, TI; Ait Bouchao, AB and Tiazite, TZ, Fig.1); when the origin of a given patient is known to the precise locality, the GPS coordinates are mentioned (Latitude; Longitude); if not, "Foum Jemaa focus" is mentioned.

| Clinical sample | Isolate | Origin, year | GPS coordinates | DNA concentration | % Leish DNA |
|---|---|---|---|---|---|
| SuSL-FJ2002 | MHOM/MO/20/FJ2002 | Tabia, 2020 | 32.02; -6.81 | 3.51 ng/ul | 26.491 |
| SuSL-FJ2003 | - | Foum Jemaa, 2020 | 31.96; -6.98 | 10.20 ng/ul | 3.559 |
| SuSL-FJ2004 | MHOM/MO/20/FJ2004 | Foum Jemaa focus, 2020 | - | 1.15 ng/ul | 2.036 |
| SuSL-FJ2005 | MHOM/MO/20/FJ2005 | Foum Jemaa, 2020 | 31.96; -6.98 | 0.73 ng/ul | 0.368 |
| SuSL-FJ2006 | - | Tabia, 2020 | 32.02, -6.81 | 2.08 ng/ul | 0.219 |
| SuSL-FJ2007 | MHOM/MO/20/FJ2007 | Tabia, 2020 | 32.02, -6.81 | 0.61 ng/ul | 3.267 |
| SuSL-FJ2008 | MHOM/MO/20/FJ2008 | Tislit, 2020 | 31.97; -6.84 | 13.4 ng/ul | 1.302 |
| SuSL-FJ2009 | - | Tislit, 2020 | 31.97; -6.84 | 8.25 ng/ul | 6.6721 |
| - | MHOM/MO/20/FJ2010 | Foum Jemaa, 2020 | 31.96; -6.98 | - | - |
| SuSL-FJ2011 | MHOM/MO/20/FJ2011 | Foum Jemaa focus, 2020 | - | 1.39 ng/ul | 0.0555 |

| | | | | | |
|---|---|---|---|---|---|
| SuSL-FJ2012 | MHOM/MO/20/FJ2012 | Ait Bouchao, 2020 | 32.03; -6.92 | 0.85 ng/ul | 2.318 |
| SuSL-FJ2013 | - | Tiazite, 2020 | 32.05; -6.85 | 7.15 ng/ul | 4.583 |
| SuSL-FJ08 | - | Foum Jemaa focus, 2018 | - | 1.35 ng/ul | 0.047 |
| SuSL-FJ26 | - | Foum Jemaa focus, 2018 | - | 2.25 ng/ul | 7.392 |
| SuSL-F28 | - | Foum Jemaa focus, 2018 | - | 7.48 ng/ul | 1.291 |
| - | MHOM/MO/16/Ltr16 | | | | |
| SuSL-IT02 | - | Imintanout, 2018 | | 1.75 ng/ul | 12.647 |
| SuSL-IT04 | - | Imintanout, 2018 | | 0.76 ng/ul | 2.342 |
| SuSL-IT08 | - | Imintanout, 2018 | | 1.31 ng/ul | 0.668 |
| - | MHOM/MO/88/M1314 | Azilal, 1988 | | | |
| - | MCAN/MO/90/M2007 | Tannant, 1990 | | | |
| - | MHOM/MO/90/M2013 | Essaouira, 1990 | | | |
| - | MHOM/MO/93/M2571 | Tannant, 1993 | | | |
| - | MHOM/MO/95/M3015 | Ouarzazate, 1995 | | | |

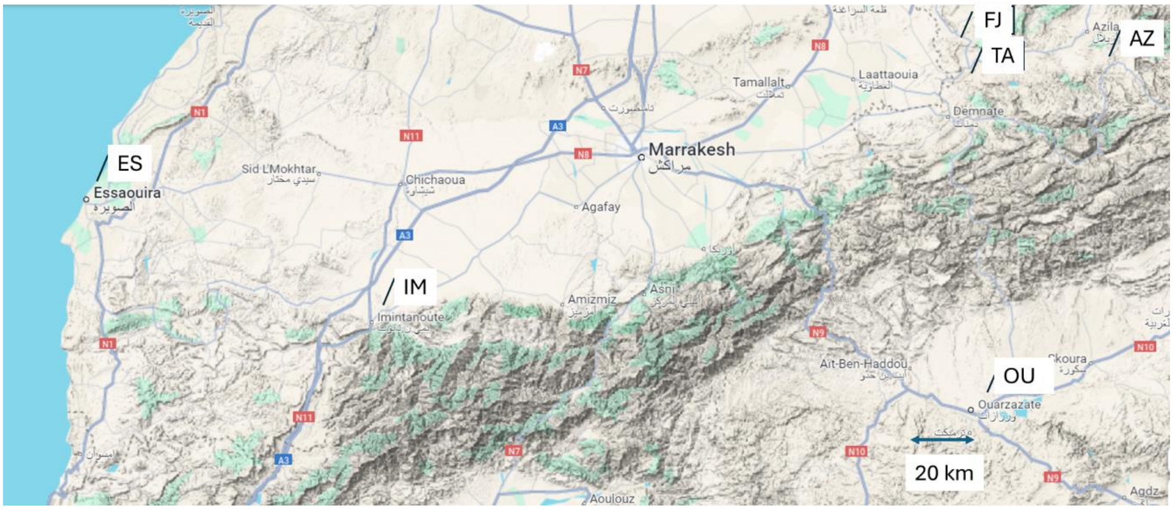


**Fig.S1.** Map showing the origin of all Moroccan samples included in the study; FJ, Foum Jemaa focus (2016-2020); AZ, Azilal (1988); TA, Tannant (1990, 1993); ES, Essaouira (1990); IM, Immintanout (2018); OU, Ouarzazate (1995). Map: © 2024 Google.

#### 2. Reference genome for analyses based on nucleotide sequences

Two reference genomes are available for *L. tropica*, respectively called TriTrypDB (L590, isolated from a human patient in Israel) and NCBI (strain CDC216-162, isolated from a human patient in Afghanistan). We assessed which one was more appropriate for mapping the sequences of our *L. tropica* samples (isolates and clinical samples). Therefore, we first proceeded to a competitive mapping of reads on *L. aethiopica* (strain L147, isolated in Ethiopia) and *L. tropica* reference genomes. This competitive mapping implies the mapping with BWA of all reads on a combined reference genome consisting of the human genome, *L. aethiopica* and *L. tropica*, and counting the number of reads mapping preferentially mapping to *L. aethiopica* versus *L. tropica* using the idxstats function of samtools. Inclusion of *L. aethiopica* was justified by the phylogenetic proximity with *L. tropica* (both belong to the same species complex). We used as controls reads of a strain of *L. aethiopica* (L100, negative), *L. tropica* (P283, positive) and a known hybrid of *L. aethiopica* and *L. tropica* (L86) [2]. When *L. tropica* NCBI reference was used, reads of all strains (including the positive control and the mix) and clinical samples mapped preferentially on the *L. aethiopica* genome (Fig. S2a). In contrast, when using the *L. tropica* TriTrypDB reference, reads of negative control mapped to *L. aethiopica* reference, those of positive control mapped to *L. tropica* reference and reads of artificial mix mapped half-half on *L. tropica* and *L. aethiopica* reference genomes (Fig. S2b).

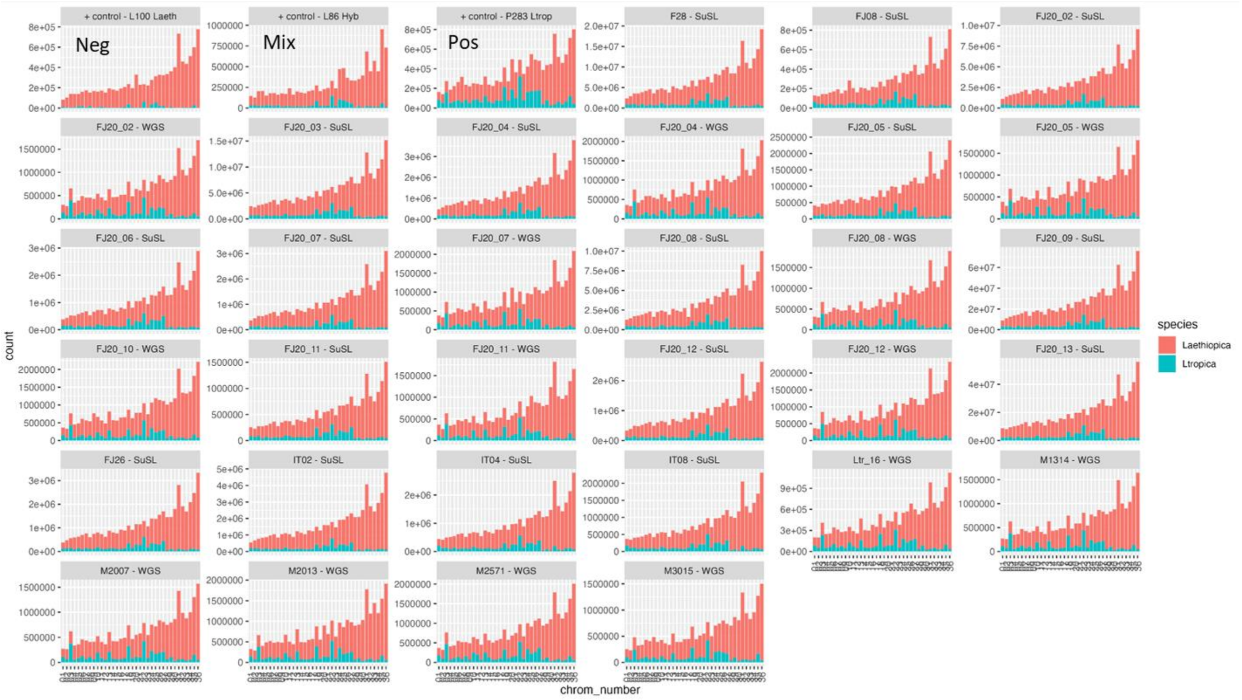


**Fig. S2a**. Reads mapped to the *L. aethiopica* L147 reference genome (represented in red) and the NCBI *L. tropica* CDC216-162 reference genome (represented in green) are displayed. The results include all genomes analyzed in this study. The X-axis represents the chromosomes, while the Y-axis indicates the number of reads mapped to each corresponding chromosome.

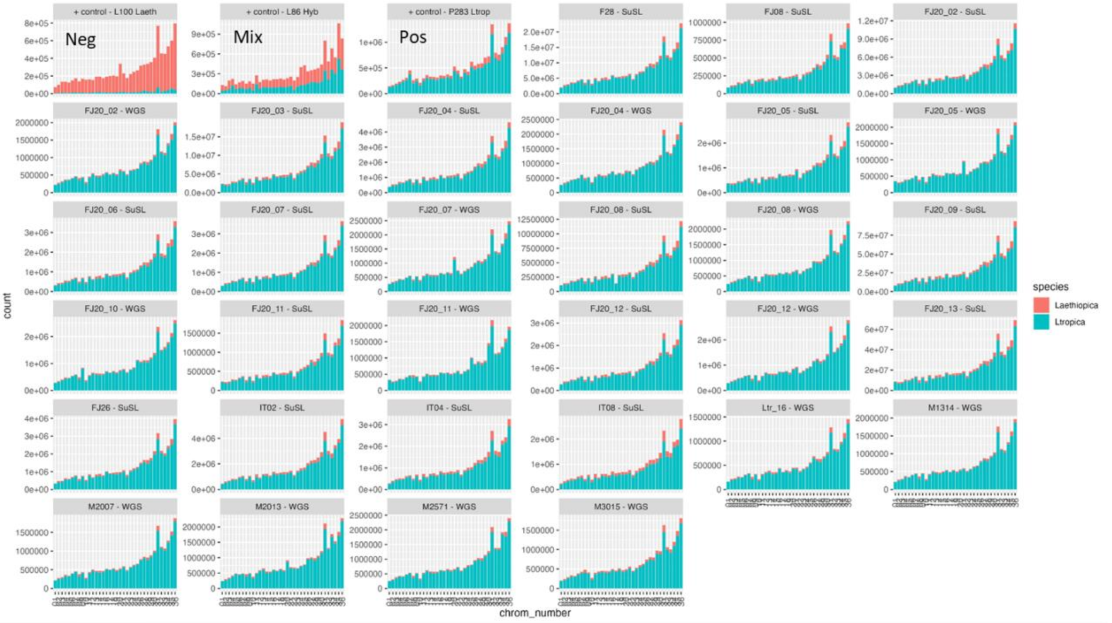


**Fig. S2b.** Reads mapped to the *L. aethiopica* L147 reference genome (represented in red) and the TriTrypDB *L. tropica* L590 reference genome (represented in green) are displayed. The results include all genomes analyzed in this study. The X-axis represents the chromosomes, while the Y-axis indicates the number of reads mapped to each corresponding chromosome.

In a second stage, we performed a phylogenetic analysis, integrating the two *L. tropica* reference genomes with those available in TriTrypDB release 64 for *L. aethiopica* L147*, L. major* strain Friedlin*,* and *L. infantum* JPCM5*,* as well as strain *L. donovani* BPK282 as published [3]. This analysis was performed on a per-chromosome basis, where corresponding chromosomes from the different strains were aligned using MAFFT and subsequently converted to a phylogenetic network. Fig.S3 shows the resulting reticulated network for 3 chromosomes, chr1, 6 and 31. The same topology was observed for each of them: *L. tropica* TriTrypDB clustered together with *L. aethiopica* and *L. major*. In contrast, *L. tropica* NCBI surprisingly clustered close together with *L. donovani* and *L. infantum,* hereby questioning the species identity of the NCBI strain. Based on competitive mapping and phylogenetics, we thus decided to use only the TriTrypDB *L. tropica* reference for further analyses of genome nucleotide sequences.

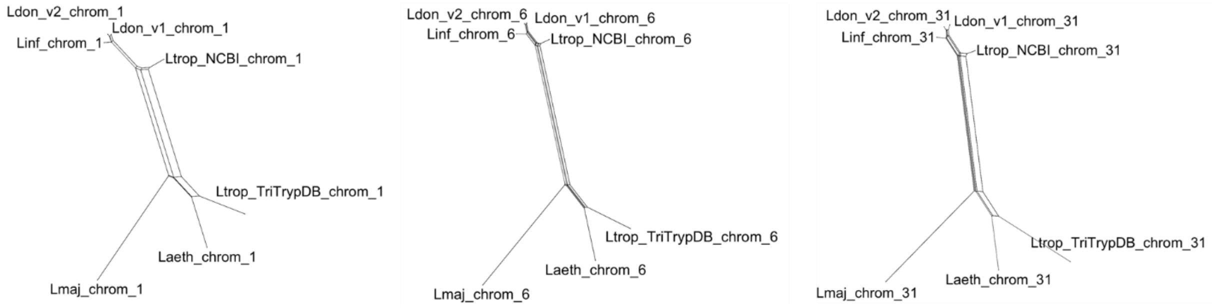


**Fig. S3.** Phylogenetic network based on nucleotide sequence of chromosomes 1, 6 and 31 in reference genomes of main *Leishmania* species of the Old World (Ltrop, *L. tropica*; Ldon, *L. donovani*; Linf, *L. infantum*; Laet, *L. aethiopica*; Lmaj, *L. major*)

**3. Reference genome for analyses of structural variations.**

For genome structure analysis (aneuploidy, local CNVs), we compared the karyotype of our *L. tropica* strains when mapping to the two *L. tropica* reference genomes (NCBI and TriTrypDB, respectively) and the closely related *L. aethiopica*. We used chr31 as a control, as this chromosome was reported by several authors as being constitutively tetrasomic (or higher), and this in all species sequenced so far, and in both promastigote and amastigote life stages. When using *L. tropica* NCBI and *L. aethiopica* references, chr 31 of Moroccan *L. tropica* strains here studied showed to be tetrasomic -as expected- while it scored as nearly disomic, when using the TriTrypDB *L.tropica* reference (Fig. S4). The latter likely resulted from the higher number of gaps in chr31 of the TriTrypDB *L. tropica* reference (Fig. S5). Accordingly, we decided to use the *L. aethiopica* reference for genome structure analyses.

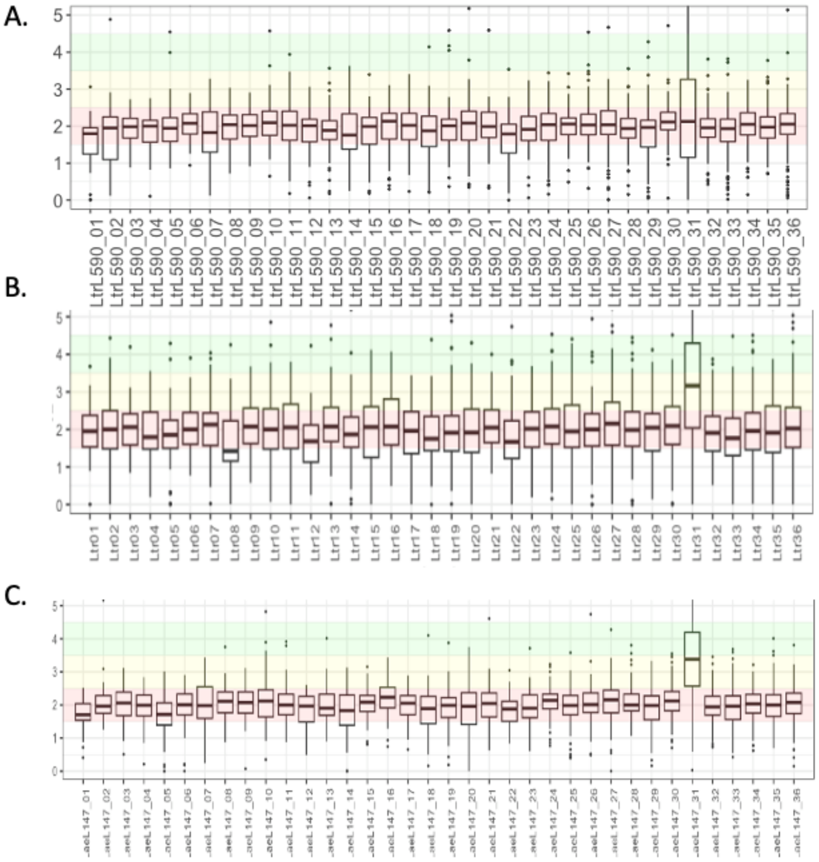

**Fig. S4.** Boxplots displaying predicted somy values for each of the 36 chromosomes, calculated from sequencing depth across 10 kb windows. The reference genomes used during read mapping are: (A) the TriTrypDB *L. tropica* reference genome, (B) the NCBI *L. tropica* reference genome, and (C) the *L. aethiopica* L147 reference genome.

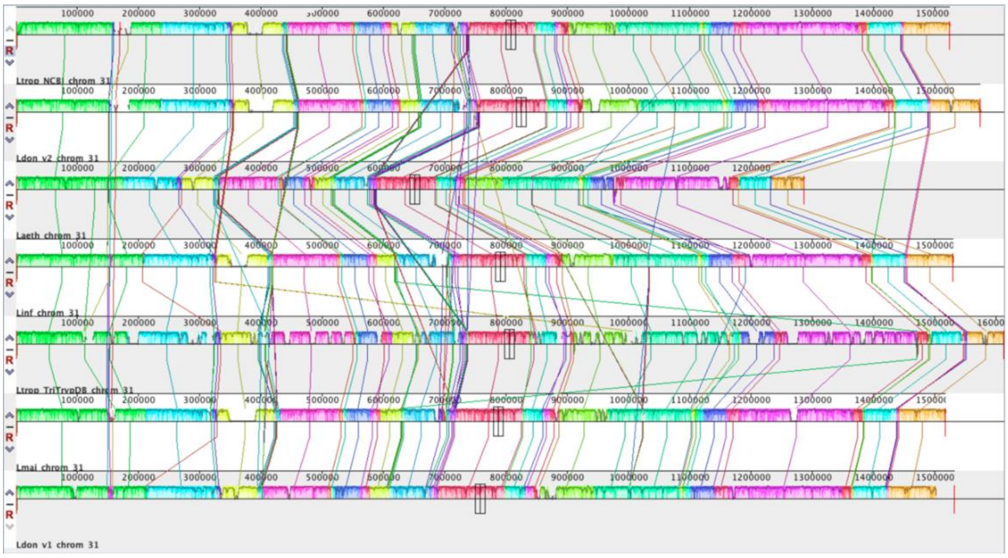


**Fig. S5.** The alignment of chromosome 31 from *Leishmania tropica* (Ltrop_NCBI) and *L. donovani* (Ldon_v1) from NCBI, along with *L. donovani* (Ldon_v2), *L. aethiopica* (Laeth), *L. infantum* (Linf), *L. tropica* (Ltrop_TriTrypDB), and *L. major* (Lmaj) from TriTrypDB. The alignment was generated using MAUVE, with homologous regions represented as locally collinear blocks (LCBs), shown as colored segments. The height of each block reflects sequence similarity, while gaps indicate regions of divergence, insertions, or deletions. The *L. tropica* TriTrypDB chromosome shows a high number of insertions and/or deletions compared to all other chromosome 31 sequences, as indicated by its sawtooth pattern, suggesting the low quality of its assembly.

**4 . Comparison of different SuSL-seq designs for capture and sequencing of *L. tropica* genome.**

We tested on *L. tropica* experimental samples two SuSL-designs that were available in the laboratory, respectively built-up with probes of *L. donovani/L. infantum* and *L. aethiopica*. We expected here that *L. aethiopica* probes would provide better results on *L. tropica* than *L. donovani*, given phylogenetic proximity of *L. aethiopica* and *L. tropica* [4]. To validate this hypothesis, we performed a benchmark experiment for which we created artificial mixes containing both *L. tropica* DNA as well as human DNA in four different concentrations of *Leishmania* DNA, i.e. 0.06%, 0.017%, 0.04% and 0.01%, called mix1 to mix4 respectively. Capturing the *L. tropica* DNA in those four artificial mixes using SureSelect was performed using two different probe designs, i.e. based on *L. donovani* and *L. aethiopica* respectively. Those results confirmed our hypothesis that genome capture of *L. tropica* showed to be possible with *L. donovani* and *L. aethiopica* using our SuSL probes, but the percentage of reads mapping to the *L. tropica* L590 genome was higher when using the SureSelect probes based on the *L. aethiopica* genome (Fig.S6a). This was confirmed running the FastQ Screen software using a genome database containing the human genome and the *L. tropica* L590 genome: for the mixes with the highest concentration (0.06% and 0.017%) we found more than 25% of the reads belonging to *L. tropica* when using the *L. aethiopica* design, while this was lower than 20% for the *L. donovani* design. For the mixes with lower amounts of *L. tropica* DNA, the difference was less clear.

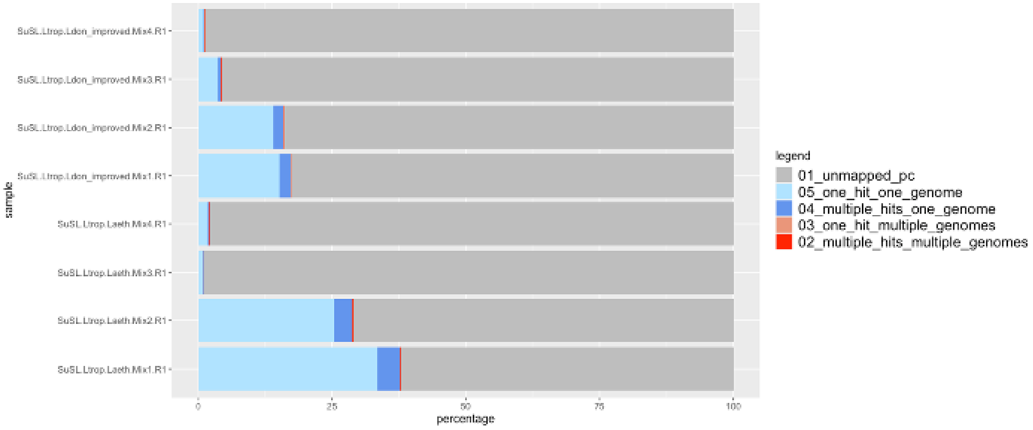


**Fig.S6a.** FastQ Screen output of four artificial mixes of *Leishmania tropica* and human DNA, containing 0.06%, 0.017%, 0.04%, and 0.01% *L. tropica* DNA, respectively. The first four bars represent the percentage of *L. tropica* reads detected using the *L. donovani* SuSL design, while the last four bars represent the percentage detected using the *L. aethopica* design. Light blue indicates reads mapping uniquely to the *L. tropica* genome, dark blue represents reads mapping to multiple locations in the *L. tropica* genome, light red shows reads mapping uniquely to both the *L. tropica* and human genomes, dark red denotes reads mapping to multiple locations in both the *L. tropica* and human genomes, and gray corresponds to reads not mapping to the *L. tropica* genome.

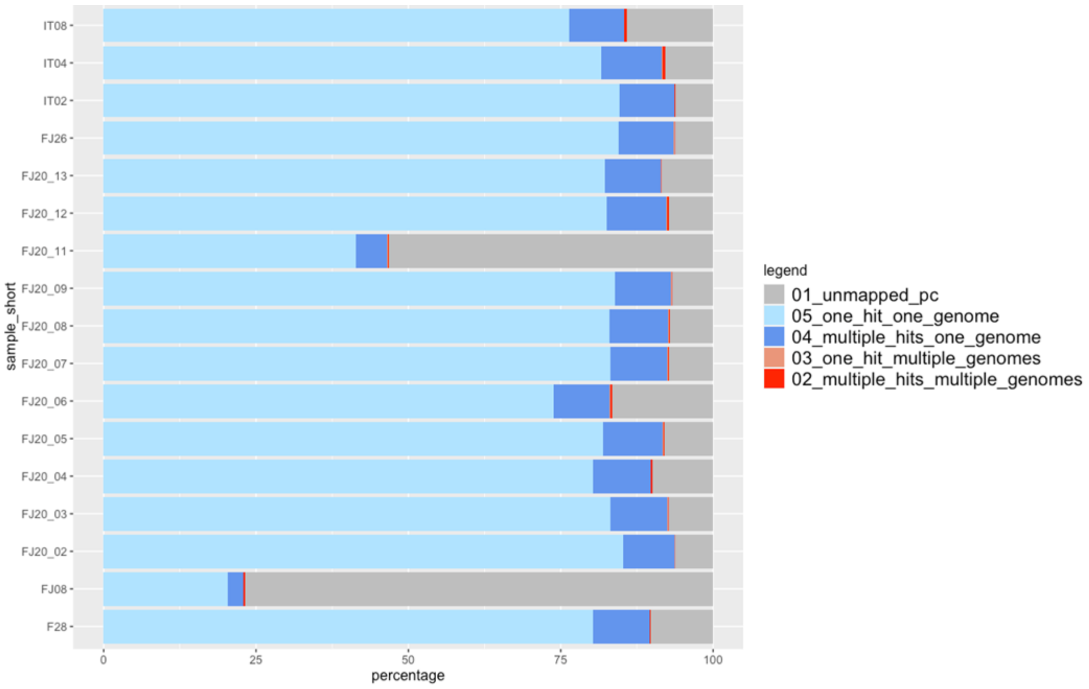


**Fig.S6b**: percentage of reads mapping to the TriTrypDB *L. tropica* reference genome among 17 clinical samples. Light blue indicates reads mapping uniquely to the *L. tropica* genome, dark blue represents reads mapping to multiple locations in the *L. tropica* genome, light red shows reads mapping uniquely to both the *L. tropica* and human genomes, dark red denotes reads mapping to multiple locations in both the *L. tropica* and human genomes, and gray corresponds to reads not mapping to the *L. tropica* genome.

##### 5. Analysis of local copy number variation in paired samples (clinical samples and derived strains)

Copy number variation (CNV) at the gene level was assessed by comparing the sequencing coverage of each gene to the median sequencing coverage of its corresponding chromosome (Fig.S7). Sequencing coverage was obtained using the samtools depth command, and CNV values were calculated using an in-house script. This script determined relative gene coverage by integrating gene annotation data from a cleaned GFF file with samtools read depth information, calculating the median coverage per gene, and normalizing it against the chromosome's median coverage. Overall, CNV estimates appeared overestimated for clinical samples analyzed with SureSelect, resulting in a median log2-normalized CNV value exceeding 0.50, whereas values were expected to peak around 0, as observed in whole-genome sequencing (WGS) data from isolates (Fig.S8). Additionally, CNV values derived from SureSelect exhibited greater spread compared to WGS-derived values, complicating the prediction of accurate CNV estimates even after correcting for the observed skewness.

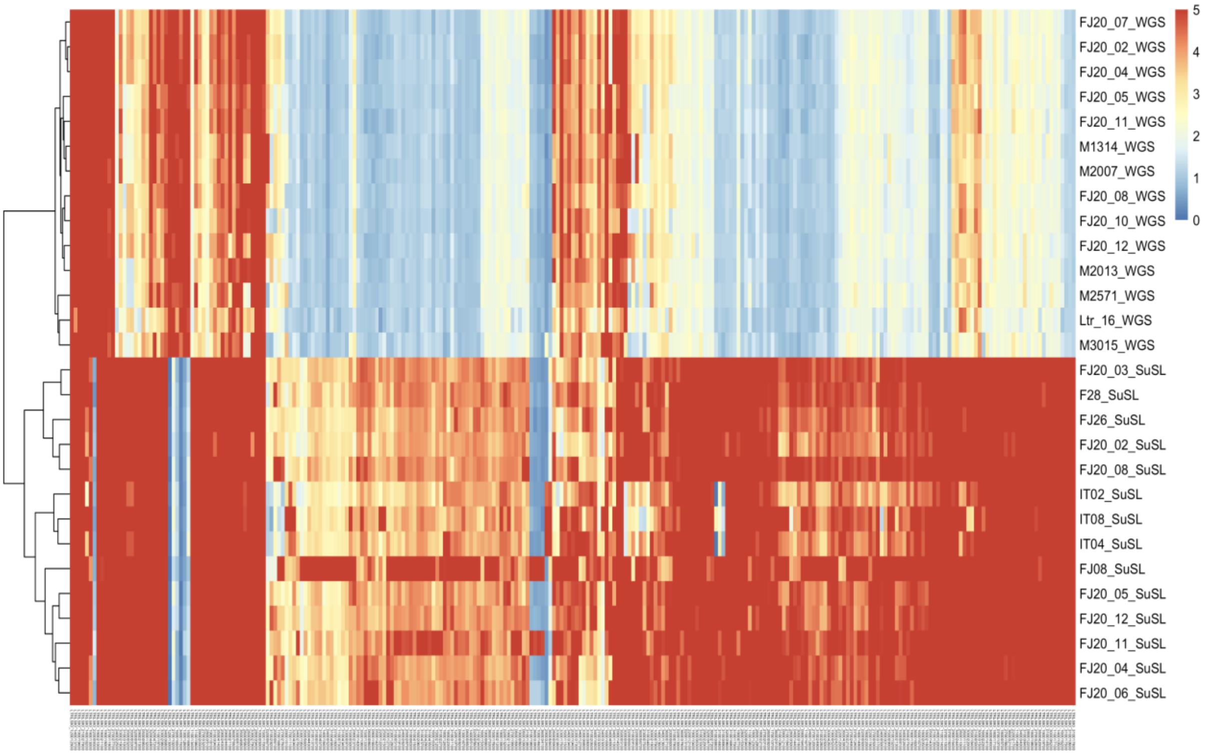

**Fig.S7** Heatmap show the log2 CNV values. In the X-axis a subset of the genes containing a CNV value higher than 10 in at least one of the samples.

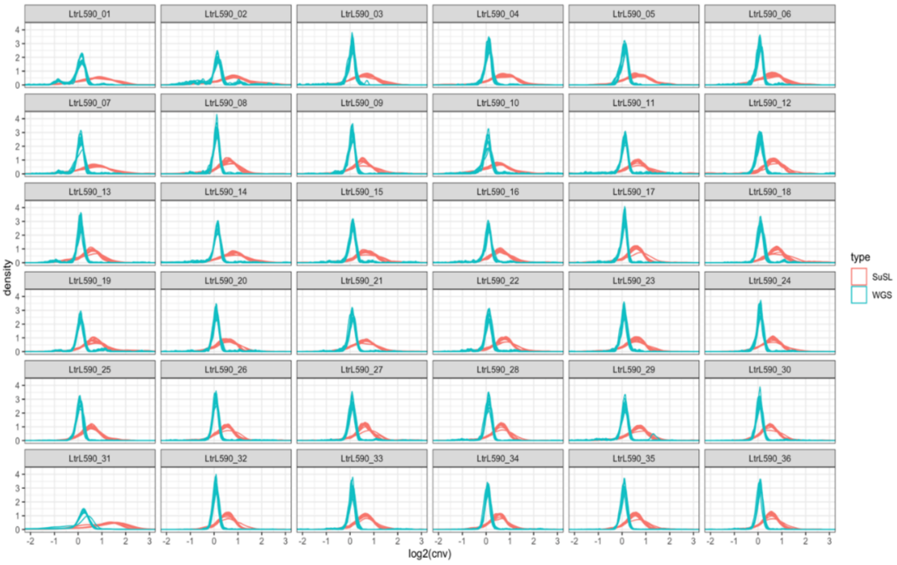


**Fig. S8** Density plot of the log2 CNV values per sequencing type (i.e. SureSelect or WGS) per sample

#### 6. Aneuploidy in paired samples

**Table S2**. Somy of the 36 chromosomes in 7 paired samples: susl, SuSL-seq of clinical samples; wgs, WGS of respective derived isolates; diff, somy difference (being biologically relevant when ≥ 0.5; pval, p-value (significant < 0.05) using Student's t-test).

| | FJ20_02 | | | | FJ20_04 | | | | FJ20_05 | | | |
|---|---|---|---|---|---|---|---|---|---|---|---|---|
| Chrom | pval | susl | wgs | diff | pval2 | susl3 | wgs4 | diff5 | pval3 | susl4 | wgs5 | diff6 |
| Ltr01 | 3,41E-01 | 1,62 | 1,89 | -0,27 | 3,45E-01 | 1,66 | 1,92 | -0,26 | 6,33E-01 | 2,58 | 2,82 | -0,24 |
| Ltr02 | 8,43E-01 | 1,88 | 2,11 | -0,23 | 9,10E-01 | 1,88 | 2,09 | -0,21 | 9,25E-01 | 1,98 | 2,14 | -0,15 |
| Ltr03 | 3,47E-01 | 1,97 | 2,20 | -0,23 | 3,45E-01 | 1,91 | 2,18 | -0,27 | 8,43E-01 | 2,03 | 2,12 | -0,09 |
| Ltr04 | 7,67E-01 | 1,91 | 2,03 | -0,12 | 7,49E-01 | 1,85 | 2,02 | -0,17 | 8,57E-01 | 1,93 | 2,00 | -0,07 |
| Ltr05 | 3,62E-01 | 1,65 | 1,98 | -0,33 | 1,38E-01 | 1,59 | 2,09 | **-0,50** | 3,74E-01 | 1,65 | 2,00 | -0,35 |
| Ltr06 | 6,86E-01 | 1,92 | 2,11 | -0,19 | 6,50E-01 | 1,95 | 2,12 | -0,17 | 7,11E-01 | 1,94 | 2,12 | -0,19 |
| Ltr07 | 8,57E-01 | 1,88 | 2,02 | -0,14 | 1,00E-01 | 1,87 | 2,19 | -0,31 | 6,90E-01 | 1,88 | 2,06 | -0,18 |
| Ltr08 | 7,40E-04 | 2,02 | 2,49 | -0,47 | 1,05E-03 | 2,00 | 2,50 | **-0,50** | 6,90E-01 | 2,06 | 2,13 | -0,07 |
| Ltr09 | 9,83E-01 | 1,99 | 2,00 | -0,01 | 8,43E-01 | 1,94 | 2,03 | -0,10 | 8,88E-01 | 2,02 | 2,04 | -0,02 |
| Ltr10 | 9,21E-01 | 2,04 | 2,02 | 0,02 | 7,71E-01 | 2,02 | 2,00 | 0,02 | 9,21E-01 | 1,96 | 2,06 | -0,11 |
| Ltr11 | 7,66E-01 | 1,92 | 1,91 | 0,01 | 5,49E-01 | 2,02 | 1,92 | 0,10 | 9,06E-01 | 1,89 | 1,95 | -0,06 |
| Ltr12 | 4,64E-01 | 1,89 | 2,03 | -0,14 | 6,90E-01 | 1,91 | 1,95 | -0,05 | 7,67E-01 | 1,94 | 1,97 | -0,04 |
| Ltr13 | 9,21E-01 | 1,86 | 1,96 | -0,10 | 6,33E-01 | 1,87 | 1,96 | -0,09 | 8,57E-01 | 1,86 | 1,98 | -0,12 |
| Ltr14 | 9,85E-01 | 1,80 | 1,94 | -0,14 | 9,76E-01 | 1,86 | 1,96 | -0,10 | 9,15E-01 | 1,84 | 1,92 | -0,08 |
| Ltr15 | 1,28E-01 | 2,05 | 2,22 | -0,16 | 9,02E-02 | 2,02 | 2,26 | -0,24 | 8,43E-01 | 1,98 | 2,03 | -0,05 |
| Ltr16 | 7,49E-01 | 2,19 | 2,14 | 0,05 | 8,43E-01 | 2,18 | 2,27 | -0,09 | 3,02E-01 | 2,23 | 2,07 | 0,16 |
| Ltr17 | 8,57E-01 | 2,01 | 2,02 | -0,01 | 1,05E-01 | 1,86 | 1,99 | -0,13 | 4,62E-01 | 2,01 | 2,09 | -0,08 |
| Ltr18 | 8,57E-01 | 1,85 | 1,92 | -0,07 | 9,40E-01 | 1,88 | 1,91 | -0,03 | 8,50E-01 | 1,77 | 1,93 | -0,16 |
| Ltr19 | 9,83E-01 | 1,96 | 1,94 | 0,02 | 9,10E-01 | 2,00 | 1,96 | 0,04 | 8,57E-01 | 1,98 | 1,91 | 0,07 |
| Ltr20 | 1,72E-01 | 1,97 | 2,28 | -0,31 | 4,64E-01 | 1,97 | 2,21 | -0,24 | 9,95E-01 | 2,05 | 1,96 | 0,09 |
| Ltr21 | 8,57E-01 | 2,06 | 2,05 | 0,01 | 8,50E-01 | 2,04 | 2,06 | -0,01 | 3,21E-05 | 2,45 | 3,06 | **-0,61** |
| Ltr22 | 6,77E-01 | 1,90 | 1,99 | -0,09 | 7,79E-01 | 1,92 | 1,97 | -0,05 | 8,50E-01 | 1,84 | 2,00 | -0,16 |
| Ltr23 | 6,72E-01 | 1,93 | 2,08 | -0,15 | 4,64E-01 | 1,95 | 2,15 | -0,19 | 8,87E-01 | 1,93 | 1,98 | -0,05 |
| Ltr24 | 1,72E-01 | 2,17 | 2,05 | 0,12 | 6,50E-01 | 2,15 | 2,22 | -0,07 | 5,99E-01 | 2,11 | 2,07 | 0,04 |
| Ltr25 | 8,37E-01 | 2,01 | 2,00 | 0,02 | 2,27E-01 | 2,01 | 2,22 | -0,21 | 9,40E-01 | 1,99 | 2,02 | -0,03 |
| Ltr26 | 8,43E-01 | 2,04 | 2,01 | 0,03 | 8,43E-01 | 2,03 | 2,01 | 0,02 | 9,95E-01 | 2,01 | 2,06 | -0,05 |
| Ltr27 | 5,20E-02 | 2,20 | 1,94 | 0,26 | 3,45E-02 | 2,20 | 1,94 | 0,26 | 1,85E-01 | 2,08 | 1,97 | 0,11 |
| Ltr28 | 4,26E-02 | 2,04 | 1,97 | 0,07 | 1,67E-02 | 2,07 | 1,95 | 0,12 | 1,91E-01 | 2,01 | 1,96 | 0,06 |
| Ltr29 | 4,86E-01 | 2,03 | 1,90 | 0,13 | 1,85E-01 | 2,05 | 1,91 | 0,14 | 6,90E-01 | 2,00 | 1,93 | 0,08 |
| Ltr30 | 1,11E-01 | 2,17 | 1,97 | 0,20 | 2,34E-02 | 2,20 | 1,97 | 0,23 | 3,62E-01 | 2,13 | 2,00 | 0,14 |
| Ltr31 | 9,21E-01 | 3,46 | 3,53 | -0,07 | 4,64E-01 | 3,67 | 3,57 | 0,10 | 8,86E-01 | 3,55 | 3,61 | -0,06 |
| Ltr32 | 2,11E-01 | 1,99 | 1,94 | 0,05 | 6,43E-02 | 2,00 | 1,95 | 0,04 | 6,70E-01 | 2,00 | 1,95 | 0,04 |
| Ltr33 | 8,50E-01 | 2,01 | 1,94 | 0,07 | 4,44E-01 | 2,05 | 1,91 | 0,14 | 8,50E-01 | 2,02 | 1,94 | 0,09 |
| Ltr34 | 8,99E-02 | 2,12 | 1,96 | 0,15 | 7,40E-04 | 2,12 | 1,93 | 0,20 | 2,34E-02 | 2,10 | 1,91 | 0,19 |
| Ltr35 | 9,06E-01 | 2,05 | 1,89 | 0,16 | 6,72E-01 | 2,06 | 1,89 | 0,17 | 8,50E-01 | 1,98 | 1,88 | 0,10 |
| Ltr36 | 1,28E-06 | 2,16 | 1,91 | 0,25 | 1,46E-08 | 2,17 | 1,92 | 0,25 | 6,67E-04 | 2,08 | 1,90 | 0,18 |

| | FJ20_07 | | | | FJ20_08 | | | | FJ20_11 | | | | FJ20_12 | | | |
|---|---|---|---|---|---|---|---|---|---|---|---|---|---|---|---|---|
| Chrom | pval4 | susl5 | wgs6 | diff7 | pval5 | susl6 | wgs7 | diff8 | pval6 | susl7 | wgs8 | diff9 | pval7 | susl8 | wgs9 | diff10 |
| Ltr01 | 2,99E-01 | 1,64 | 1,93 | -0,29 | 8,57E-01 | 1,78 | 1,95 | -0,17 | 1,85E-01 | 2,53 | 2,93 | -0,40 | 6,86E-01 | 1,73 | 1,87 | -0,14 |
| Ltr02 | 9,90E-01 | 1,92 | 2,08 | -0,16 | 8,88E-01 | 1,88 | 2,07 | -0,19 | 9,21E-01 | 1,94 | 2,06 | -0,12 | 8,50E-01 | 2,00 | 1,99 | 0,00 |
| Ltr03 | 6,50E-01 | 1,95 | 2,18 | -0,23 | 4,98E-01 | 1,99 | 2,17 | -0,18 | 9,21E-01 | 1,91 | 2,13 | -0,23 | 8,50E-01 | 2,05 | 2,17 | -0,11 |
| Ltr04 | 6,72E-01 | 1,93 | 2,02 | -0,09 | 8,50E-01 | 1,90 | 2,02 | -0,12 | 8,37E-01 | 1,91 | 2,04 | -0,13 | 9,31E-01 | 1,94 | 1,99 | -0,05 |
| Ltr05 | 5,36E-01 | 1,63 | 1,88 | -0,25 | 3,62E-01 | 1,66 | 2,01 | -0,35 | 7,67E-01 | 1,67 | 1,89 | -0,22 | 2,99E-01 | 1,66 | 2,04 | -0,38 |
| Ltr06 | 7,66E-01 | 1,98 | 2,12 | -0,14 | 8,08E-01 | 1,93 | 2,15 | -0,22 | 6,50E-01 | 1,94 | 2,14 | -0,20 | 4,79E-01 | 1,99 | 2,24 | -0,24 |
| Ltr07 | 8,43E-01 | 1,80 | 2,05 | -0,24 | 8,67E-01 | 1,90 | 1,99 | -0,09 | 8,43E-01 | 1,95 | 2,05 | -0,10 | 9,15E-01 | 1,86 | 1,97 | -0,11 |
| Ltr08 | 6,89E-01 | 2,02 | 2,08 | -0,06 | 3,45E-01 | 2,01 | 2,20 | -0,19 | 8,79E-05 | 1,93 | 2,81 | **-0,87** | 9,21E-01 | 2,10 | 2,03 | 0,08 |
| Ltr09 | 8,50E-01 | 2,01 | 2,03 | -0,01 | 9,32E-01 | 2,00 | 2,06 | -0,06 | 8,36E-01 | 2,00 | 2,00 | 0,00 | 9,15E-01 | 1,98 | 2,05 | -0,06 |
| Ltr10 | 4,02E-01 | 2,00 | 2,33 | -0,33 | 8,43E-01 | 2,00 | 2,05 | -0,05 | 8,00E-01 | 1,99 | 1,98 | 0,01 | 9,30E-01 | 2,00 | 2,04 | -0,04 |
| Ltr11 | 7,66E-01 | 1,95 | 1,93 | 0,02 | 5,15E-01 | 1,96 | 1,94 | 0,02 | 3,45E-01 | 2,00 | 1,90 | 0,10 | 3,69E-01 | 1,97 | 1,84 | 0,14 |
| Ltr12 | 7,66E-01 | 1,89 | 1,93 | -0,04 | 7,28E-01 | 1,91 | 1,98 | -0,07 | 8,43E-01 | 1,76 | 1,97 | -0,22 | 8,43E-01 | 1,92 | 1,97 | -0,06 |
| Ltr13 | 9,12E-01 | 1,86 | 2,00 | -0,14 | 9,21E-01 | 1,90 | 2,01 | -0,11 | 7,11E-01 | 1,89 | 1,97 | -0,07 | 9,21E-01 | 1,87 | 2,01 | -0,14 |
| Ltr14 | 9,36E-01 | 1,83 | 1,95 | -0,11 | 9,30E-01 | 1,86 | 1,95 | -0,09 | 9,21E-01 | 1,81 | 1,94 | -0,13 | 9,36E-01 | 1,86 | 1,95 | -0,09 |
| Ltr15 | 8,19E-01 | 1,95 | 2,01 | -0,06 | 8,57E-01 | 2,05 | 2,04 | 0,00 | 9,54E-01 | 2,05 | 2,03 | 0,02 | 9,21E-01 | 1,99 | 2,02 | -0,03 |
| Ltr16 | 6,93E-02 | 2,24 | 2,05 | 0,19 | 5,58E-02 | 2,20 | 2,03 | 0,17 | 1,85E-01 | 2,21 | 2,08 | 0,13 | 1,22E-01 | 2,23 | 2,07 | 0,16 |
| Ltr17 | 8,50E-01 | 2,00 | 1,98 | 0,01 | 3,07E-31 | 1,08 | 2,01 | **-0,93** | 6,50E-01 | 2,04 | 2,08 | -0,03 | 4,52E-01 | 2,00 | 2,08 | -0,08 |
| Ltr18 | 8,57E-01 | 1,83 | 1,96 | -0,13 | 9,32E-01 | 1,89 | 1,94 | -0,05 | 9,54E-01 | 1,84 | 1,91 | -0,07 | 8,50E-01 | 1,85 | 1,95 | -0,11 |
| Ltr19 | 9,30E-01 | 1,99 | 1,96 | 0,03 | 8,57E-01 | 2,01 | 1,99 | 0,02 | 7,67E-01 | 1,99 | 1,93 | 0,06 | 8,43E-01 | 2,02 | 1,92 | 0,10 |
| Ltr20 | 2,44E-08 | 2,02 | 3,50 | **-1,48** | 5,73E-01 | 1,98 | 2,10 | -0,12 | 8,54E-01 | 1,95 | 2,01 | -0,06 | 3,69E-01 | 2,01 | 2,20 | -0,19 |
| Ltr21 | 9,21E-01 | 2,02 | 2,07 | -0,05 | 8,50E-01 | 2,08 | 2,07 | 0,01 | 7,31E-01 | 2,04 | 2,04 | 0,00 | 6,99E-02 | 2,07 | 2,33 | -0,26 |
| Ltr22 | 8,57E-01 | 1,93 | 1,98 | -0,05 | 8,57E-01 | 1,94 | 1,99 | -0,05 | 9,30E-01 | 1,83 | 1,94 | -0,10 | 8,50E-01 | 1,93 | 1,91 | 0,02 |
| Ltr23 | 9,95E-01 | 1,92 | 1,97 | -0,05 | 4,57E-01 | 1,95 | 1,96 | -0,02 | 6,86E-01 | 1,98 | 1,98 | 0,00 | 3,21E-01 | 1,95 | 1,89 | 0,06 |
| Ltr24 | 3,62E-01 | 2,15 | 2,06 | 0,09 | 1,53E-01 | 2,17 | 2,07 | 0,10 | 4,82E-02 | 2,15 | 2,04 | 0,11 | 1,00E-01 | 2,12 | 2,01 | 0,11 |
| Ltr25 | 8,43E-01 | 2,02 | 2,11 | -0,09 | 8,26E-01 | 2,01 | 2,03 | -0,02 | 6,07E-12 | 2,12 | 2,96 | **-0,84** | 9,96E-01 | 2,00 | 2,04 | -0,04 |
| Ltr26 | 8,43E-01 | 2,04 | 2,01 | 0,02 | 8,57E-01 | 2,06 | 2,16 | -0,10 | 9,32E-01 | 2,00 | 2,01 | -0,02 | 8,50E-01 | 2,05 | 2,10 | -0,06 |
| Ltr27 | 1,85E-01 | 2,14 | 1,98 | 0,16 | 4,82E-02 | 2,23 | 1,97 | 0,26 | 1,67E-02 | 2,28 | 1,99 | 0,29 | 6,50E-01 | 2,14 | 2,02 | 0,12 |
| Ltr28 | 1,01E-01 | 2,04 | 2,00 | 0,04 | 2,77E-02 | 2,08 | 1,98 | 0,09 | 2,82E-02 | 2,06 | 1,96 | 0,10 | 2,82E-02 | 2,03 | 1,93 | 0,10 |
| Ltr29 | 6,20E-01 | 2,03 | 1,95 | 0,07 | 4,86E-01 | 2,04 | 1,94 | 0,11 | 2,87E-01 | 2,04 | 1,90 | 0,15 | 1,30E-01 | 2,04 | 1,85 | 0,19 |
| Ltr30 | 1,85E-01 | 2,17 | 2,02 | 0,16 | 3,45E-02 | 2,20 | 1,99 | 0,21 | 2,29E-08 | 2,21 | 2,71 | -0,50 | 5,20E-02 | 2,15 | 1,97 | 0,19 |
| Ltr31 | 8,43E-01 | 3,60 | 3,61 | -0,01 | 6,90E-01 | 3,61 | 3,56 | 0,05 | 8,19E-03 | 3,76 | 4,33 | **-0,57** | 9,69E-01 | 3,54 | 3,66 | -0,13 |
| Ltr32 | 3,69E-01 | 2,00 | 1,98 | 0,03 | 1,29E-01 | 2,00 | 1,97 | 0,03 | 9,02E-02 | 2,00 | 1,94 | 0,06 | 1,19E-02 | 1,96 | 1,89 | 0,07 |
| Ltr33 | 8,50E-01 | 2,01 | 1,94 | 0,08 | 4,62E-01 | 2,07 | 1,93 | 0,14 | 9,54E-01 | 2,03 | 2,02 | 0,01 | 2,15E-01 | 2,02 | 2,13 | -0,12 |
| Ltr34 | 1,67E-02 | 2,14 | 1,98 | 0,17 | 4,75E-03 | 2,17 | 1,96 | 0,21 | 3,46E-04 | 2,06 | 1,91 | 0,15 | 9,34E-03 | 2,09 | 1,91 | 0,18 |
| Ltr35 | 9,83E-01 | 2,02 | 1,95 | 0,07 | 7,75E-01 | 2,06 | 1,94 | 0,12 | 1,07E-01 | 2,07 | 1,87 | 0,20 | 5,11E-01 | 1,99 | 1,84 | 0,15 |
| Ltr36 | 6,67E-04 | 2,15 | 1,96 | 0,18 | 1,91E-06 | 2,20 | 1,96 | 0,24 | 3,09E-04 | 2,15 | 1,89 | 0,26 | 2,44E-03 | 2,10 | 1,95 | 0,15 |

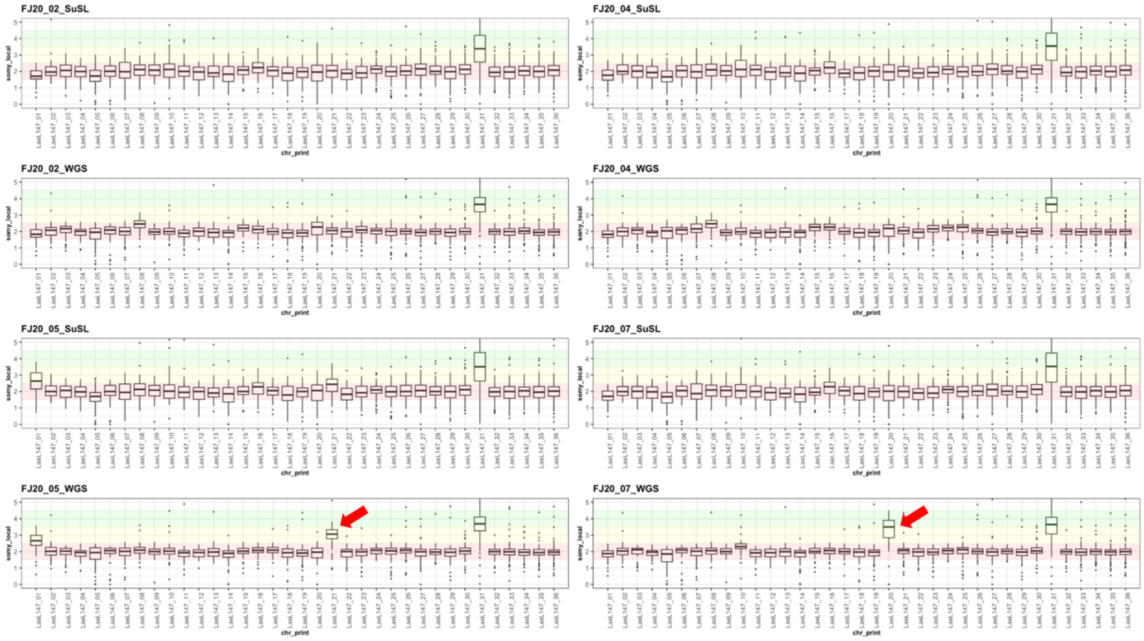


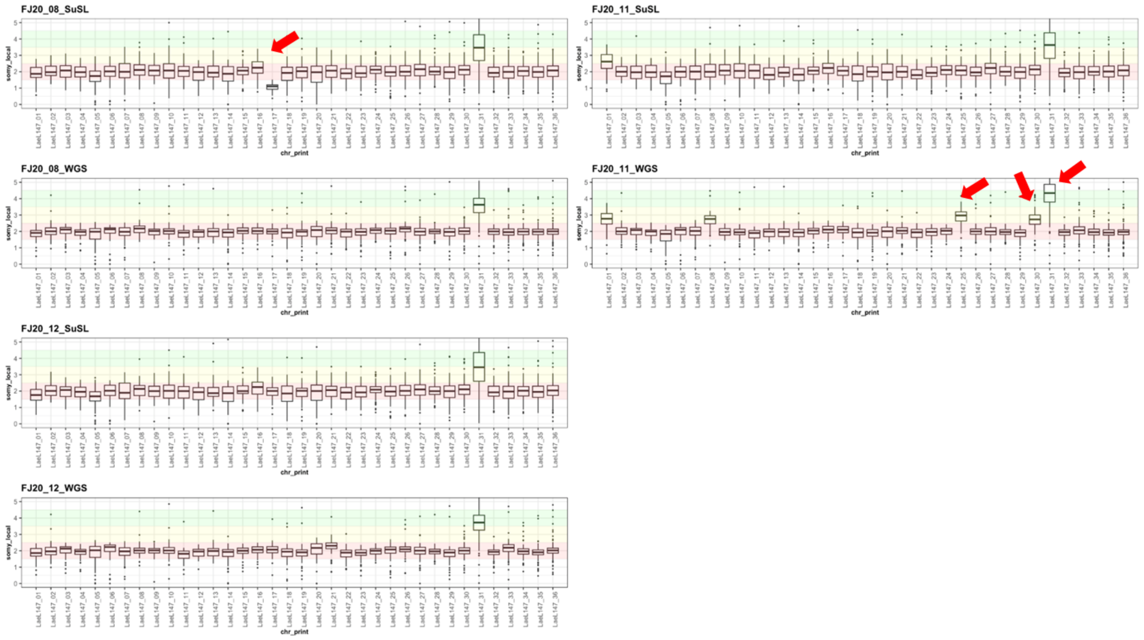


**Fig. S9.** Karyotypes of paired samples (for each patient, SuSL-seq, clinical sample and WGS, derived isolate); the arrow marks the chromosomes showing a significant change in somy between paired samples (see criteria in legend of Table 1).

#### 7. Phylogenomics

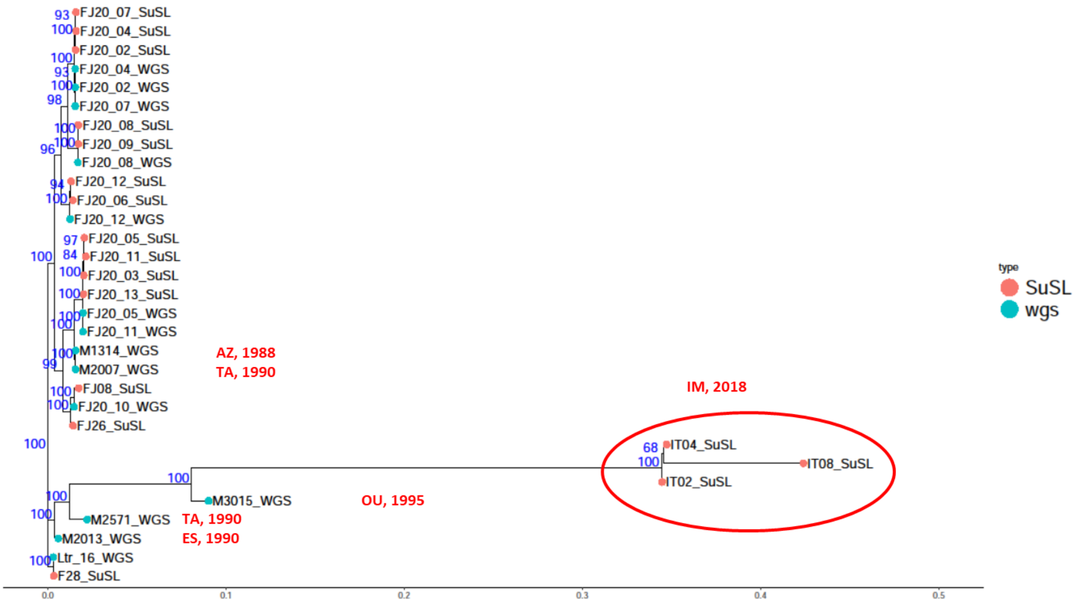


**Fig.S10.** Rooted phylogenetic tree based on genome-wide bi-allelic SNPs using RAxML, with an indication of the relevant boostrap values in blue. All samples originate from the Foum Jemaa focus except when indicated in red (TA, Tannant; AZ, Azilal; ES, Essaouira; OU, Ouarzazate; IM, Imintanout) together with the year of sampling.

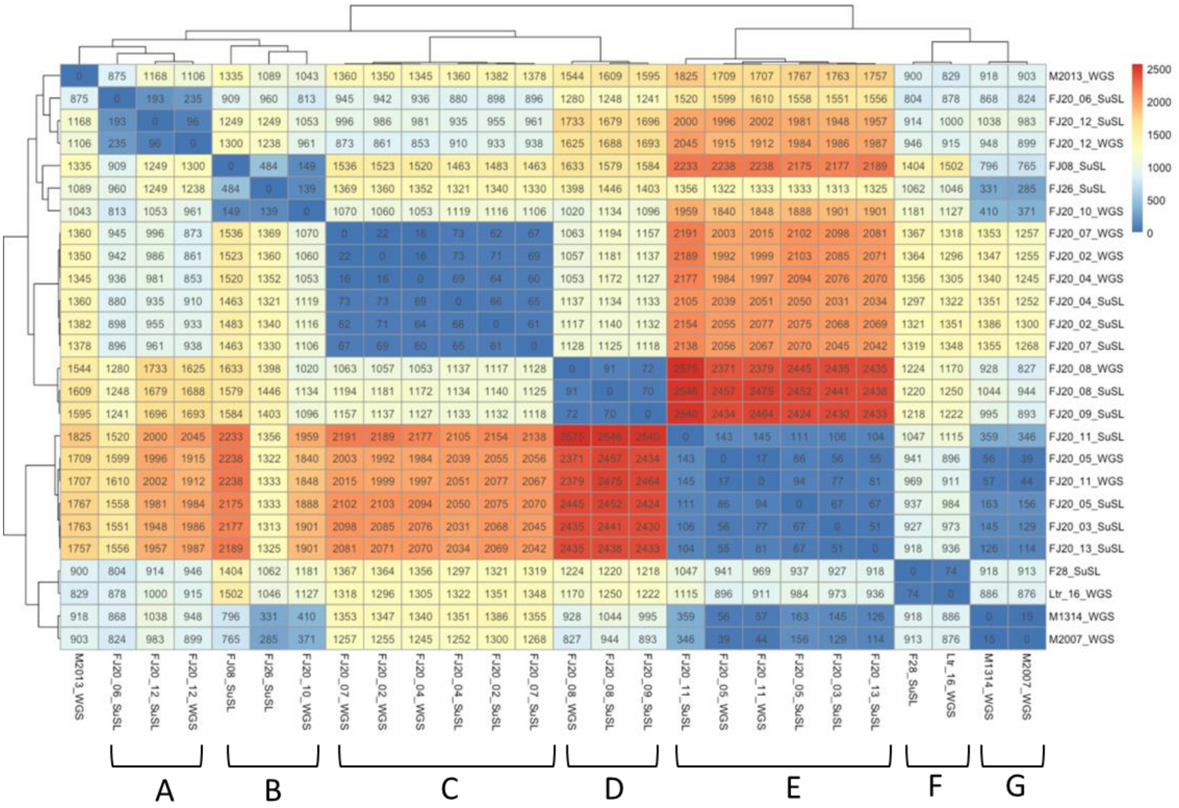


**Fig.S11.** Heatmap showing the number of homozygous SNPs overlapping between each pair of strains. A, 96 to 235 SNPs, average 174; B, 139 to 484 SNPs, average, 257; C, 16 to 73 SNPs, average 59; D, 70 to 91 SNPs, average 177; E, 17 to 145 SNPs, average 84; F, 74 SNPs; G, 15 SNPs.

#### 8. Recombination

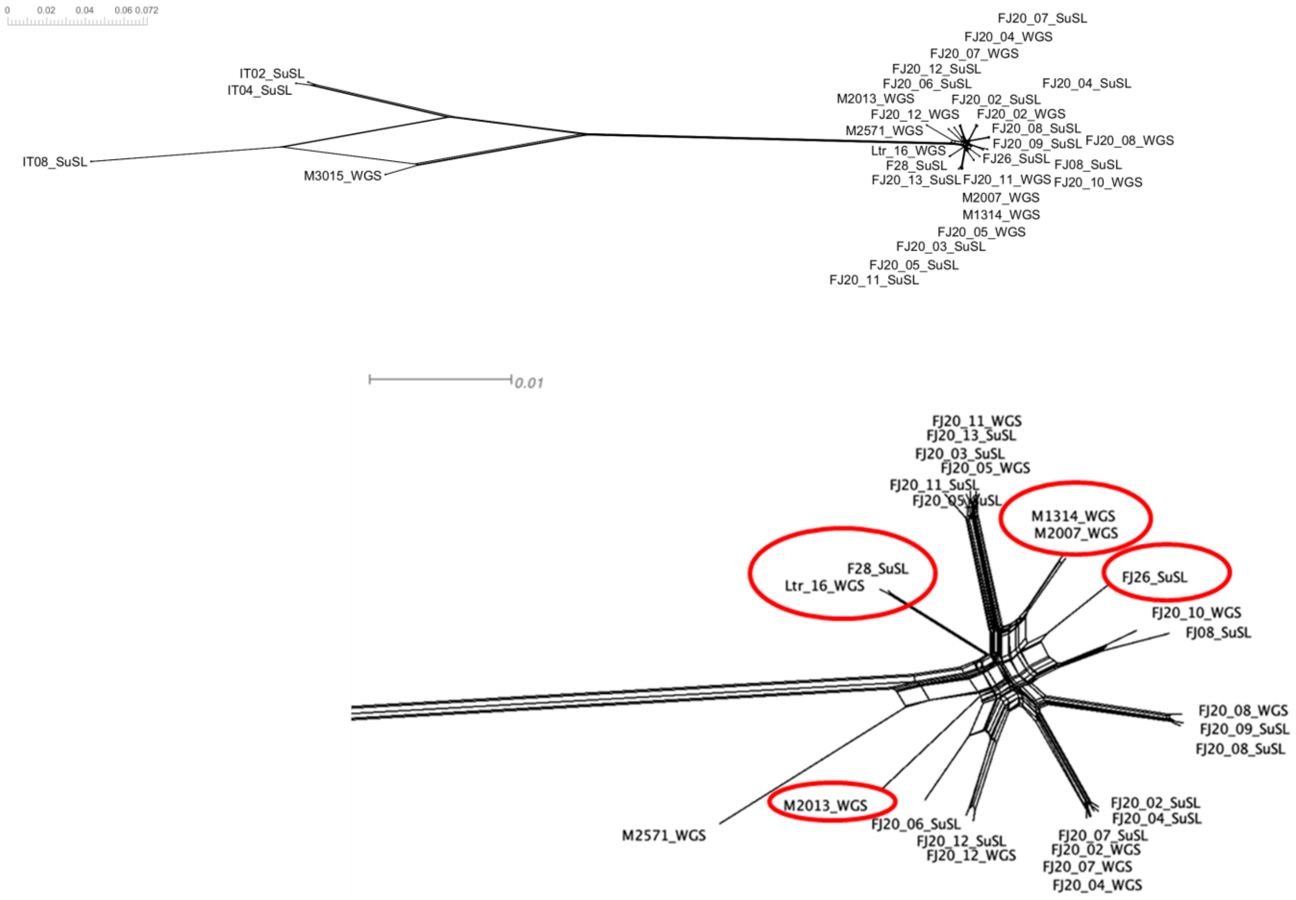


**Fig.S12.** Phylogenetic network created based on 463,062 bi-allelic SNPs using the NeighborNet algorithm as implemented in SplitsTree4. The upper panel shows all samples while the lower panel shows a zoom on samples from the Foum Jemaa focus. The six samples highlighted in red oval correspond to samples for which admixture analysis (Fig. 4) showed clear patterns of mixed ancestry. The seventh admixed sample (M3015) was an outlier in phylogenetic analysis shown in Fig.S10 and upper panel.

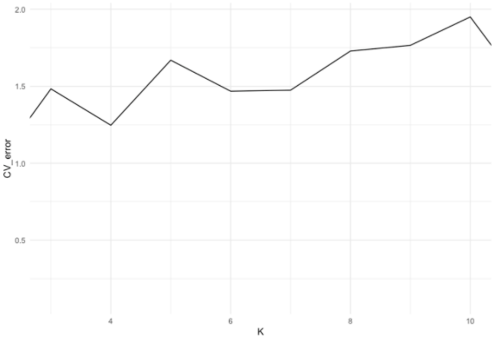


**Fig.S13.** Cross validation error as extracted from the ADMIXTURE log files for each tested value of the number of populations (K).

Available from: https://bmcgenomics.biomedcentral.com/articles/10.1186/s12864-019-6126-y
